# Multimodal profiling establishes ovarian fibrosis as a measurable and targetable hallmark of human reproductive aging

**DOI:** 10.64898/2026.09.04.749443

**Authors:** Lydia Hughes, Emily Zaniker-Gomez, Pooja R. Devrukhkar, Subhasri Biswas, Alexis Trofimchuk, Tomiris Atazhanova, Joan K. Riley, Anna Kleinhans, Man Zhang, Signe Holm Nielsen, Morten Karsdal, Michael B. Stout, Elnur Babayev, Francesca E. Duncan

## Abstract

Ovarian aging underpins infertility, systemic morbidity, and mortality in women. Stromal fibrosis is implicated in ovarian aging but has not been defined in the human ovary in situ. We integrated shear wave elastography (SWE), extracellular matrix (ECM) neoepitope fingerprinting, and single-cell RNA sequencing of follicular fluid aspirates within the same individual to comprehensively profile ovarian fibrosis. In an age-stratified cohort (≤33 and ≥37 years; N=32), age predicted ovarian stiffness, while mean stiffness was independently associated with reduced oocyte yield and follicular efficiency. ECM profiling revealed a dynamic fibrotic index favoring formation over degradation products. Single-cell transcriptomics demonstrated an age-associated fibroinflammatory stromal program enriched in pro-fibrotic pathways with stromal–immune crosstalk. In a larger validation cohort (25–45 years; N=100), age was similarly associated with ovarian stiffness. Thus, ovarian fibrosis is a quantifiable hallmark of human reproductive aging and represents a robust potential non-invasive biomarker and target for therapeutic intervention.

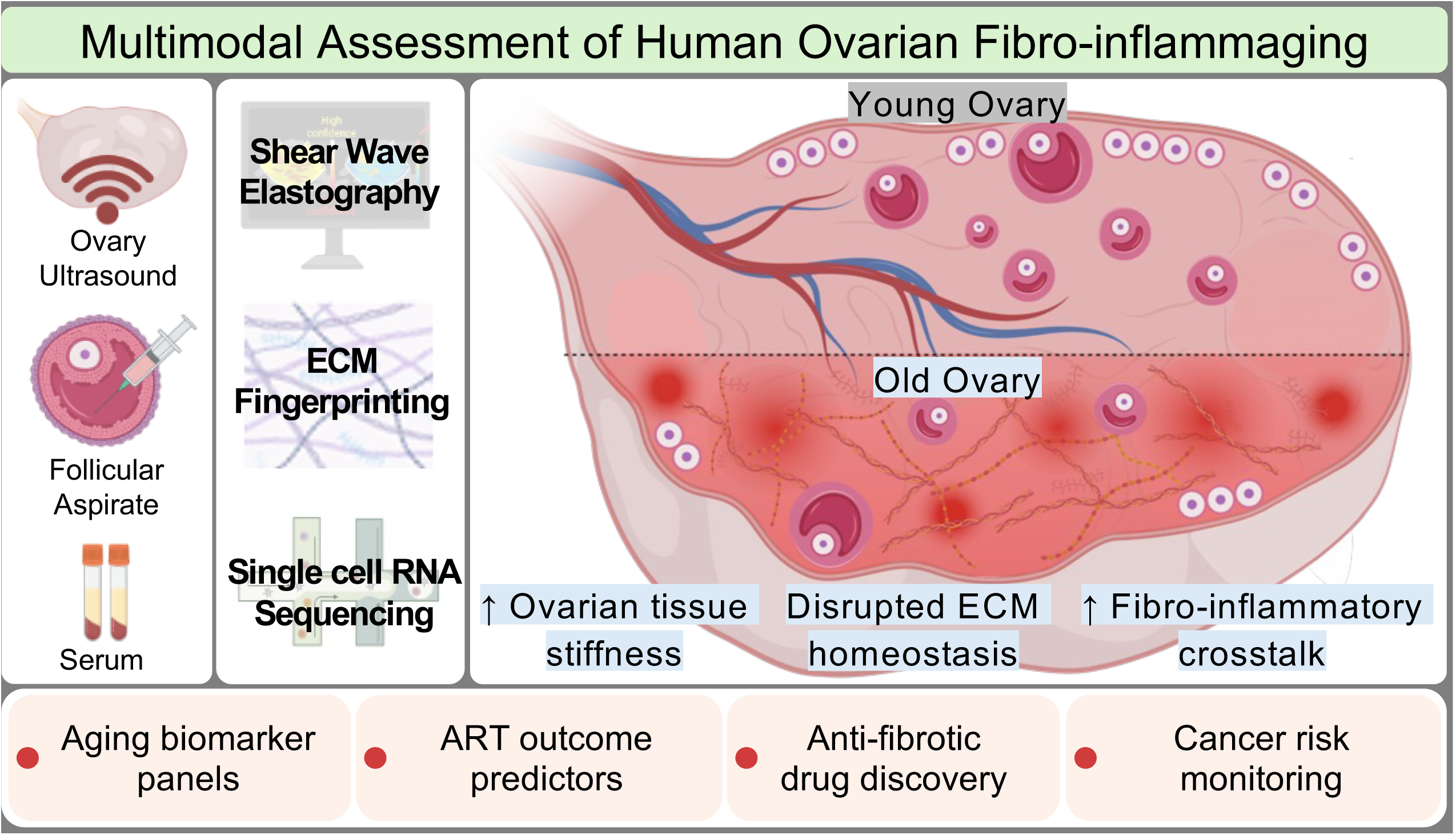

## INTRODUCTION

Ovarian aging represents the earliest and most clinically consequential manifestation of biological aging in women, preceding dysfunction of most other organ systems by decades. Reproductive aging is characterized by the progressive decline in oocyte quantity and quality, directly increasing rates of infertility and miscarriage while driving global reliance on Assisted Reproductive Technologies (ART).^1,2^ Beyond reproduction, ovarian aging has profound systemic consequences due to decreased follicular estrogen production contributing to bone loss, cardiometabolic dysfunction, and neurocognitive decline.^3,4^ Earlier age of menopause is independently associated with increased risk of osteoporosis^5–7^, composite cardiovascular disease^8,9^, dementia^10^, and all-cause mortality.^11^ Notably, while human lifespan has increased substantially over time, the age at menopause has remained relatively constant, prolonging the duration of post-reproductive life and extending exposure to estrogen deficiency.^4^ Given these far-reaching clinical and societal implications, there is an urgent need to define the mechanisms underlying ovarian aging to develop therapeutic strategies.

Fibrosis is a conserved hallmark of human aging—a process in which progressive extracellular matrix (ECM) accumulation, crosslinking, and stiffening disrupts normal tissue architecture, function, and drives systemic morbidity. This fibroinflammatory remodeling underlies pathologic and age-associated functional decline in the kidney^12^, liver^13^, lung^14^, and heart.^15,16^ In mice, the ovary also becomes fibro-inflammatory with age, characterized by altered immune cell composition and fibroblast gene expression, excess collagen I and III deposition, loss of hyaluronan, and broader matrisome dysregulation.^17–26^ ^1,18,20,25–28^. This age-dependent stromal fibrosis is associated with a quantifiable increase in ovarian tissue stiffness and has biological consequences on follicle development and function, oocyte quality, and ovulation capacity.^18,22,24,25,29–34^ While age-associated ovarian fibrosis appears to be conserved in human, this analysis has been primarily limited to histological analyses of fixed tissues which fails to capture the complex biomechanical and molecular dynamics of this fibroproliferative condition.^18,23,26,35^

This study fundamentally advances our mechanistic understanding of human ovarian aging by using a multimodal approach to measure biomechanical and molecular indices of ovarian fibrosis within the same individual in an age-stratified cohort undergoing planned fertility preservation or assisted reproduction for non-infertile indications (≤33 or ≥37 years old, N=32). We first used transvaginal shear wave elastography (SWE), an ultrasound-based technology, to quantify ovarian tissue stiffness in situ. We then performed ECM neoepitope fingerprinting to detect specific matrix protein formation and degradation products that were released into follicular fluid and serum, enabling a dynamic assessment of active ovarian and systemic ECM turnover, respectively.^36,37^ Finally, single-cell RNA sequencing (scRNA-seq) was performed on follicular fluid cellular aspirates collected during oocyte retrieval to characterize the cellular landscape and age-related transcriptomic changes within the periovulatory ovarian microenvironment. We demonstrate that human ovarian aging is defined by quantifiable biomechanical stiffening, pro-fibrotic ECM remodeling across biological compartments, and fibro-inflammatory stromal transcriptional interactions. Importantly, this ovarian stiffness was related to direct clinical endpoints given that increased stiffness was associated with reduced oocyte yield and follicular efficiency independent of age, anti-Müllerian Hormone (AMH), and body mass index (BMI) in the context of ART. Furthermore, the age-dependent increase in human ovarian stiffness was validated prospectively in an independent continuous age cohort (24–45 years old, N=100), demonstrating the broad significance of these findings. Thus, ovarian fibrosis has high potential for biomarker development and therapeutic targeting as a modifiable determinant of reproductive longevity.

## RESULTS

### Human ovarian stiffness increases with age in situ

To define human ovarian fibrosis in situ, we developed an integrated multimodal pipeline combining transvaginal SWE in the early follicular phase, cycle days 2-3, of the menstrual cycle (in two independent but complementary cohorts), targeted ECM neoepitope profiling of paired serum and follicular fluid, scRNA-seq of antral follicle cellular aspirates, and analysis of ART outcomes **(Fig. 1a).** Cohort 1 was age-stratified into reproductively young (≤33 years; n=16) and reproductively old (≥37 years; n=16) participants undergoing ART. To minimize confounding from other infertility etiologies, the younger group was restricted to participants with non-female infertility indications, while the older group included participants with non-female infertility indications, diminished ovarian reserve, or unexplained infertility, the latter two being attributable to ovarian aging (Cohort 1 Demographics; **Supplementary Table 1).** As expected, participants in the reproductively young and old groups exhibited significant age-related differences in ovarian reserve markers, including decreased AMH and antral follicle count (AFC) **(Supplementary Table 1)**. There was also a significant difference in BMI between the two groups, with higher BMI in the older cohort, underscoring the importance of controlling for BMI in multivariate analyses **(Supplementary Table 1)**. In Cohort 1 SWE was performed during the follicular phase of the menstrual cycle (days 2-3) **(Fig. 1a-ii).** Serum and follicular fluid pooled from 3 dominant follicles per ovary were collected at the time of oocyte retrieval. ECM neoepitope profiling was performed on serum and follicular fluid from a subset of n=10 per group, and scRNA-seq was performed on follicular fluid cellular aspirate from n=5 per group **(Fig. 1a-iii).** Cohort 2 was comprised of participants across the reproductive age continuum (N=100; 24–45 years) and were used to prospectively validate the ovarian stiffness findings from Cohort 1, with SWE also performed on days 2-3 of the menstrual cycle **(right panel: Fig. 1a-i)**.

**Fig 1.**
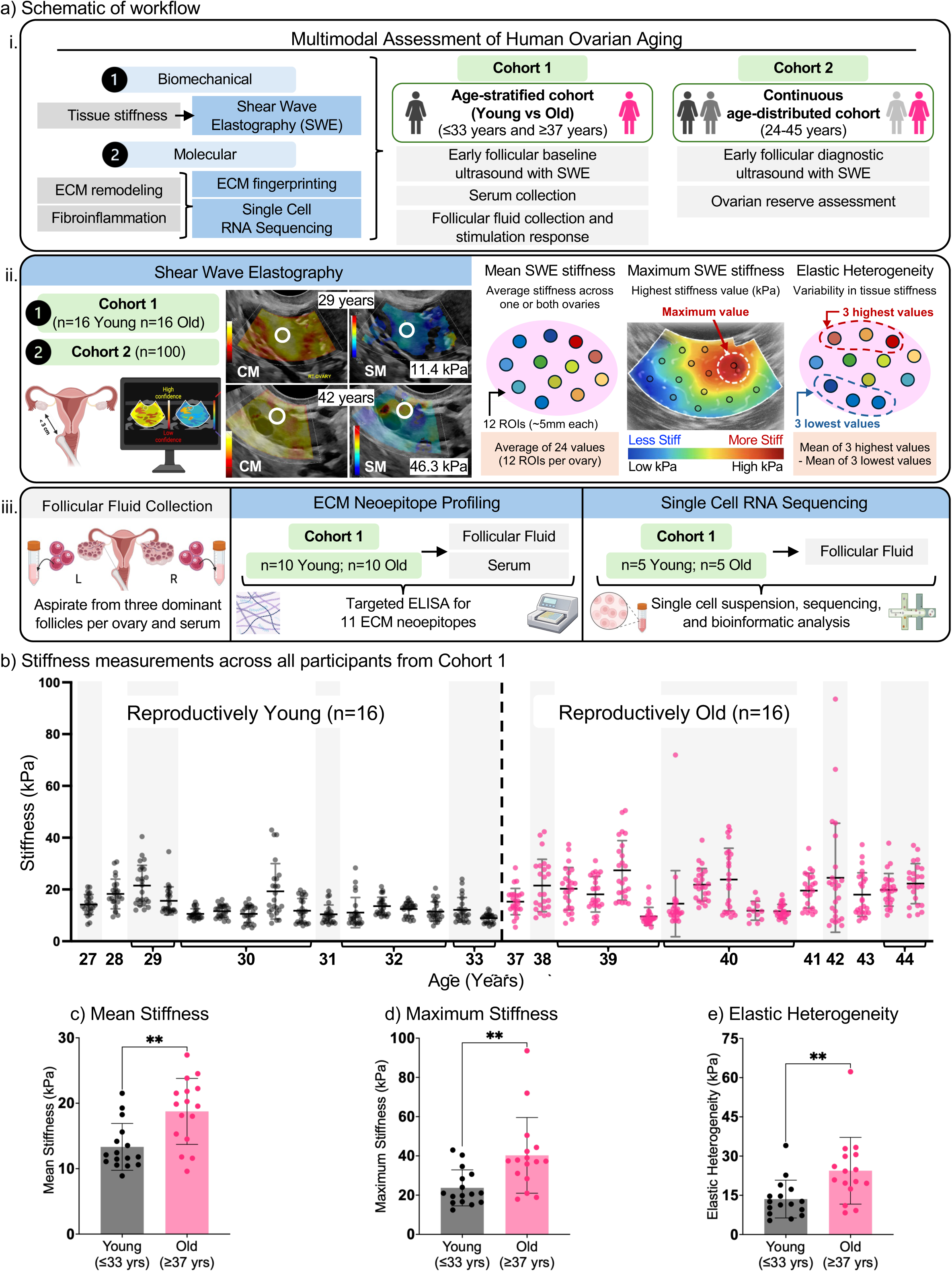
Human ovarian stiffness increases with reproductive age. **a, i.** Schematic of multimodal study design (left) performed on age-stratified Cohort 1 and continuous age-distributed Cohort 2 (right), **ii.** Transvaginal shearwave elastography (SWE) performed in early follicular phase (menstrual cycle day 2-3) for Cohort 1 and 2; representative SWE images of a confidence map with corresponding color map of max stiffness (kPa) from reproductively young (29 years) and old (42 years) participants from Cohort 1 (left). Three ovarian stiffness parameters derived per participant (right)**. iii,** Outline of molecular analyses performed in subset of Cohort 1 participants**. b,** Individual SWE stromal stiffness measurements for all Cohort 1 participants plotted by age, stratified into reproductively young (left, *n=*16) and reproductively old (right, *n=* 16) groups. Each data point represents a single ROI measurement (*n=*24 measurements per participant); black lines are mean ± SD. **c-e**, Bar graphs of mean stiffness **(c),** maximum stiffness **(d),** and elastic heterogeneity **(e)** between reproductively young versus old participants in Cohort 1; Bars represent mean ± SD. **p<0.01 by unpaired t-test.

To assess ovarian stiffness, SWE measurements were acquired across a maximum of 12 overlapping, high-confidence regions of interest (∼5 mm each) per ovary throughout the stroma, yielding up to 24 stiffness values per participant (representative ovary; **Supplementary Fig. 1**). From these measurements, we analyzed three complementary biomechanical parameters that have demonstrated clinical relevance in non-reproductive medical contexts: mean stiffness (average across all ROIs), maximum stiffness (highest ROI value), and elastic heterogeneity (EH) (mean of the three highest values minus the mean of the three lowest), capturing the magnitude and spatial distribution of ovarian stromal stiffness **(Fig. 1a-ii)**.^38–42^ Ovarian stiffness measurements ranged from 5.2 to 43.0 kPa in reproductively young participants and from 5.4 to 93.6 kPa in reproductive old. Intra-individual variability in stiffness measurements, assessed as the standard deviation of repeated measurements per ovary, was greater in reproductively old participants than reproductively young participants (median SD 3.8 vs. 7.3 kPa; Mann-Whitney U, p = 0.003). This pattern persisted after normalizing for elevated mean stiffness in the older group (coefficient of variation 32.0% vs. 37.0%; p = 0.05; **Fig. 1b**). All three parameters of ovarian stiffness were consistently higher in the older group, including mean stiffness (13.3 ± 3.6 vs 18.8 ± 5.0 kPa, p <0.01), maximum stiffness (23.7 ± 9.2 vs 40.3 ± 19.3 kPa, p <0.01), and EH (13.6 ± 7.2 vs 24.4 ± 12.8 kPa, p <0.01) **(Fig.1c-e).** These data indicate that the human ovary becomes stiffer wholistically with age and also exhibits areas of increased focal rigidity. After adjusting for AMH levels, a hormone which is an indirect measure of follicle quantity and ovarian reserve, and BMI, age ≥37 remained significantly associated with increased stiffness: mean (β1 = 7.8, p<0.001), maximum (β1 =21.6, p<0.01), and EH (β1 = 14.6, p<0.01) **(Table 1)**.^43–46^ Independent of ovarian reserve and body composition, older age was associated with an ∼8-22 kPa increase in various stiffness parameters. BMI was modestly (∼0.7 and 1.8 kPa, p<0.05) inversely associated with mean stiffness and EH (**Table 1).**

**Table 1.** Association of age with ovarian stiffness in Cohort 1. Models were fit for the ovarian stiffness parameter (kPa) as a dependent variable, with age category (reproductively old ≥37 versus young ≤33 years), AMH (ng/mL), and BMI (kg/m²) as covariates. Estimates represent adjusted differences in stiffness (kPa).

| Mean Stiffness |  |  |  |
| --- | --- | --- | --- |
| Characteristic | Estimate | 95% CI | p-value |
| Age (Young/Old) | 7.80 | 4.1 to 11.5 | <0.001* |
| AMH | 0.73 | -0.4 to 1.8 | 0.18 |
| BMI | -0.70 | -1.4 to 0.02 | 0.04* |
| Elastic Heterogeneity |  |  |  |
| Characteristic | Estimate | 95% CI | p-value |
| Age (Young/Old) | 14.62 | 5.8 to 23.4 | <0.01* |
| AMH | 0.16 | -2.5 to 2.8 | 0.90 |
| BMI | -1.78 | -3.4 to -0.16 | 0.03* |
| Maximum Stiffness |  |  |  |
| Characteristic | Estimate | 95% CI | p-value |
| Age (Young/Old) | 21.58 | 8.6 to 34.6 | <0.01* |
| AMH | 0.20 | -3.7 to 4.1 | 0.92 |
| BMI | -2.37 | -4.8 to 0.03 | 0.053 |

### Ovarian stiffness is inversely correlated with clinical outcomes in ART cycles

Because increased rigidity of the ovarian microenvironment can negatively impact follicle development and egg quality in animal models, we next assessed whether increased human ovarian stiffness would restrict oocyte yield and follicular efficiency (FORT and FOI) as the most proximal clinical measures of follicle development in ART cycles.^18,22,30,47–49^ Higher mean ovarian stiffness was independently associated with reduced total and mature oocyte yield from ART cycles (**Table 2**). Moreover, ovarian stiffness was inversely associated with parameters of follicular efficiency after adjusting for age, AMH, and BMI **(Table 2).** Follicular output rate (FORT) is a measure of follicle response to stimulation^50,51^ and is defined as the ratio of the number of pre-ovulatory follicles (≥16 mm) on the day of ovulation trigger injection relative to the baseline AFC at the start of stimulation. This parameter exhibited a negative association with mean ovarian stiffness (β1 = −2.84, p = 0.046). Follicular output index (FOI) is a measure of oocyte yield in response to stimulation^52,53^ and is calculated as the ratio of total oocytes retrieved relative to the baseline AFC. This parameter also exhibited a negative association with mean ovarian stiffness (β1 = −5.47, p = 0.03). The relationships between ovarian stiffness, age, and oocyte outcomes during ART indicate that the biomechanical signature of ovarian aging captures an aspect of ovarian function not fully explained by chronological age or circulating ovarian reserve markers.

**Table 2.**
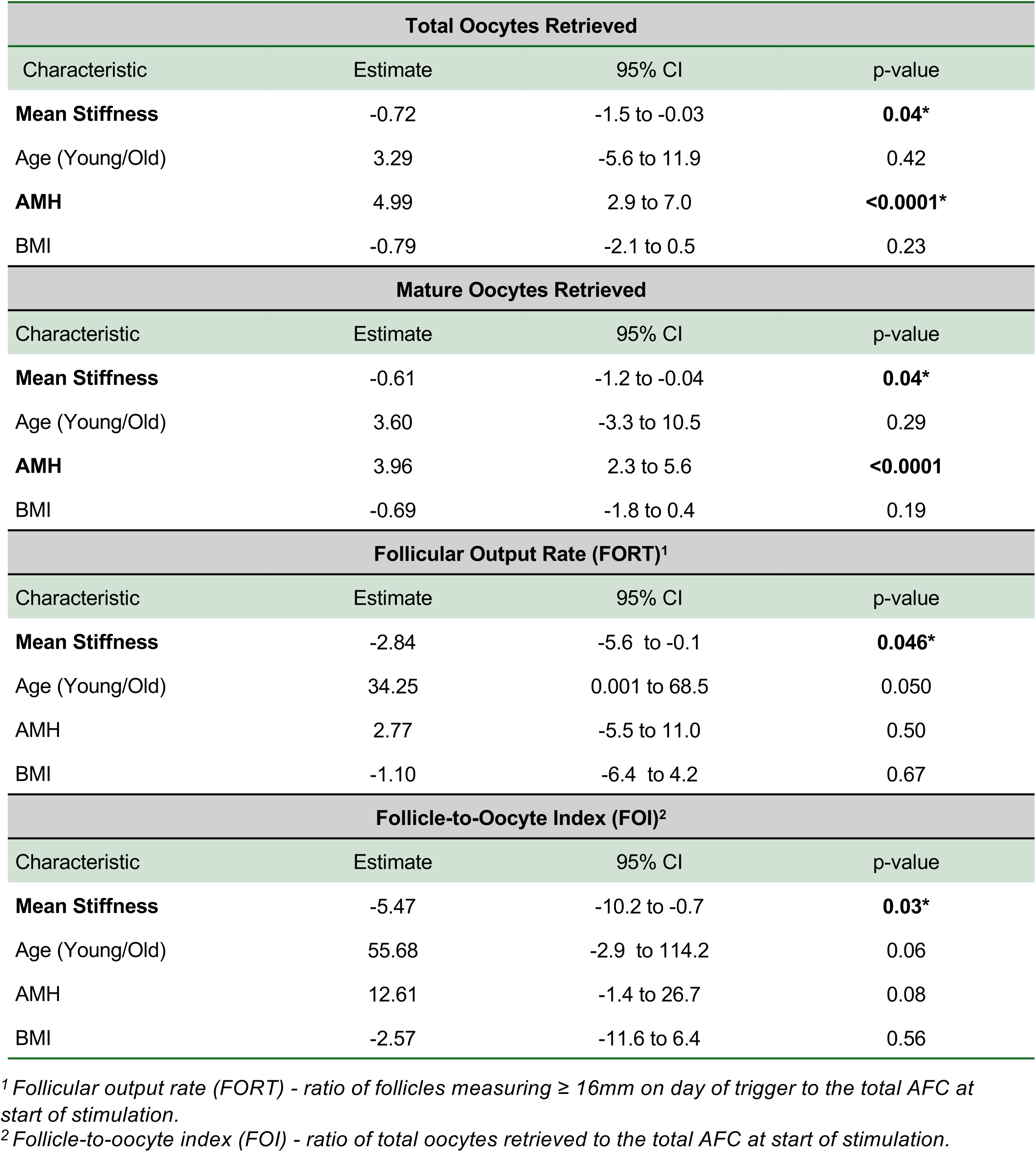
Association of ovarian stiffness with ART outcomes. Models were fit for the oocyte or follicular efficiency outcome as the dependent variable with mean ovarian stiffness (kPa), AMH (ng/mL), and BMI (kg/m²) as covariates. Estimates represent adjusted differences in oocyte or follicular efficiency parameter.

### ECM neoepitope profiling reveals a novel fibro-proliferative signature of ovarian aging

To determine whether the biomechanical stiffness of the ovary is reflective of fibrosis, we investigated dynamic ovarian fibrotic remodeling across paired follicular fluid and serum, representing ovarian and systemic biofluids, respectively. Because fibrosis reflects dysregulation of both matrix formation and degradation, we applied ECM neoepitope profiling to capture this dynamic remodeling process **(Fig. 2a).** This ELISA-based ProteinFingerprint^TM^ technology detects specific peptide fragments, termed neoepitopes, that are released into biofluids during enzymatic cleavage of collagens and proteoglycans (matrix degradation), alongside pro-peptides shed during procollagen processing (matrix formation), thereby simultaneously quantifying active ECM turnover.^54^ Participant demographics of the subset of Cohort 1 used for ECM neoepitope profiling (n= 20 total) were representative of the entire cohort (n=32 total) **(Fig. 2b).** We developed a custom panel of 11 neoepitopes based on proteins that have been implicated in the aging ovary (Nordic Bioscience A/S, Herlev, Denmark; **Supplementary Table 3**).^35,55,56^ Overall, we observed an enrichment of collagen formation epitopes relative to degradation epitopes in both serum and follicular fluid, with PRO-C8 (collagen VIII, Vastatin) exhibiting the largest age-related increase in fold change in follicular fluid (1.8-fold) **(Fig. 2c).** PRO-C8 and C6M (MMP-degraded collagen VI) were the neoepitopes with the most pronounced age-related differences (**Fig. 2d** and **Supplementary Fig. 2**). Mean PRO-C8 levels in follicular fluid (0.4±0.3 vs 0.7±0.5 ng/mL, p= 0.16) trended higher in the older group indicating increased collagen VIII formation with age (**Top panel; Fig. 2d**). In contrast, follicular fluid C6M was significantly lower in the older group (8.4±1.2 vs 7.2±1.1 ng/mL, p= 0.03) indicating reduced collagen VI degradation with age **(Bottom panel; Fig. 2d**). Intraindividual neoepitope levels in the follicular fluid directly correlated to levels in the serum (**Fig. 2e**). Together these findings are reflective of a net fibroproliferative ovarian milieu with age.

**Fig 2.**
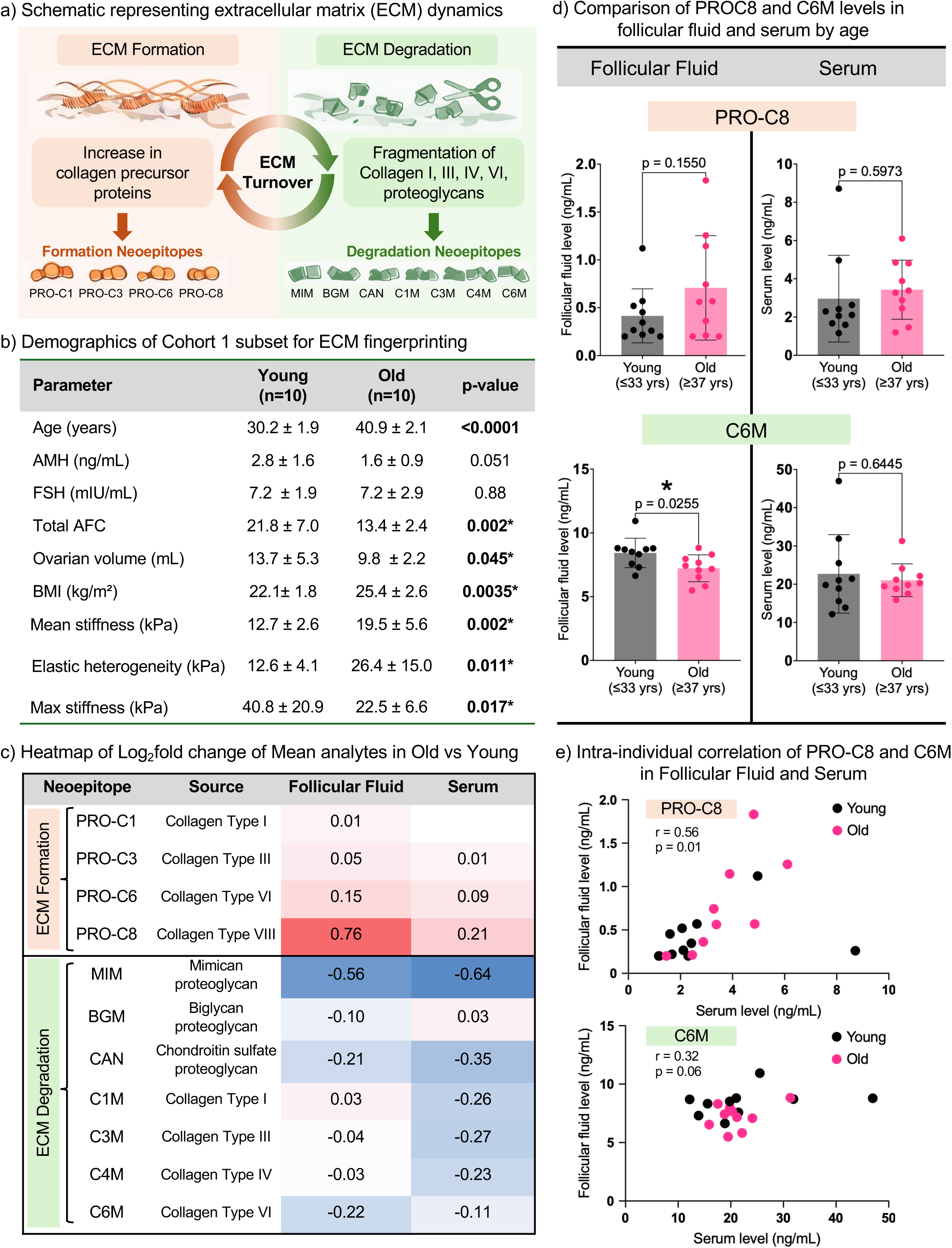
ECM fingerprinting reveals disrupted matrix homeostasis in the aged ovarian microenvironment. **a,** Diagram of dynamic nature of extracellular matrix (ECM) remodeling with target formation and degradation neoepitopes. **b,** Demographics table of subset of Cohort 1 participants for ECM fingerprinting analysis, (*n=*10 young, *n=*10 old per group). Table values in mean ± SD, comparisons by unpaired t-test. **c,** Heatmap depicting log₂ fold change of mean neoepitope levels in reproductively old relative to young participants across serum and follicular fluid compartments. Values >0 indicate enrichment with age (red scale); values <0 indicate down shift with age (blue scale). **d,** Comparison of select neoepitopes levels, PRO-C8 (top) and C6M (bottom), by age group in follicular fluid and serum. Bar graphs represent mean ± SD; comparisons by unpaired t-test, *p <0.05. **e,** Scatter plot of intra-individual correlations of PRO-C8 (top; positive, r= 0.56, *p= 0.01) and C6M (bottom; positive, r= 0.32, p= 0.06) in paired serum and follicular fluid.

### Ovarian stromal cells orchestrate a fibroinflammatory transcriptional program with reproductive aging

To resolve the molecular phenotype of human ovarian aging at the single-cell level, we performed single cell RNA sequencing of follicular fluid cellular aspirates collected during oocyte retrieval in a subset of 5 reproductively young (30.2 ± 2.2 years old) and 5 reproductively old (41.0 ± 2.0 years old) participants from Cohort 1 that had similar demographic and ovarian stiffness parameters to the broader cohort (**Fig. 3a, 3b**). Unsupervised clustering identified twelve distinct cell populations that were identified based on established marker genes^57–62^ (**Fig. 3c, 3d**). The predominant cell types in both age groups were immune cells (neutrophils, macrophages, and T cell lineages) and granulosa and granulosa/theca cells, consistent with the cell source of periovulatory follicles following gonadotropin stimulation (**Fig. 3e**). Overall cellular composition was similar between groups, but aspirates from reproductively young participants contained a higher mean proportion of stromal cells (**Fig. 3e**), which may reflect a relative increase in size of the fibrotic stromal tissue in older ovaries.

**Fig 3.**
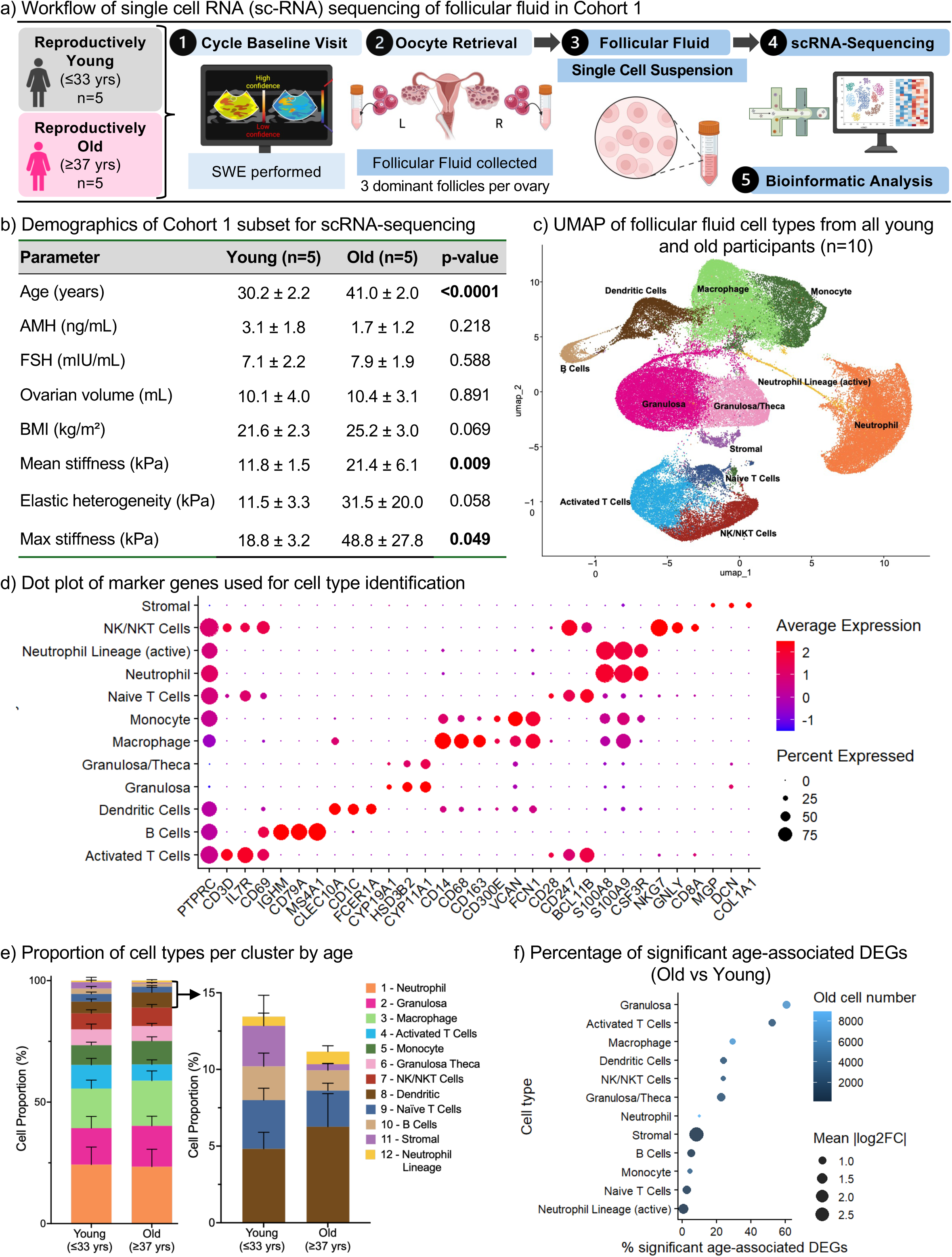
Single-cell transcriptomics of follicular fluid reveals widespread age-associated transcriptional reprogramming. **a,** Schematic of single cell RNA sequencing (scRNA-seq) workflow. **b,** Demographics table of subset of Cohort 1 participants for scRNA-seq analysis, (*n=*5 young and *n=*5 old participants). Table values are in mean ± SD; group comparisons by unpaired t-test **c,** UMAP plots of all cells from combined young and old participants (*n=*10 total participants; total cells = 92,509) identifying twelve distinct populations. **(d)** Dot plot of canonical lineage marker genes used for cluster annotation. Dot size indicates the percentage of cells expressing each gene; color reflects scaled average expression (low to high reflected as blue to red). **e,** Stacked bar plots reflecting proportional cellular composition per age group (left) and expanded view of less abundant clusters (right). Error bars represent SEM. **f,** Bubble plot displaying percentage of significant age-associated differentially expressed genes (DEGs) in old compared to young group (adjusted p ≤0.0001) per cluster (x-axis), with bubble size representing mean absolute log2 fold change and color indicating total cell number in the reproductively old group (scale low to high = dark to light blue).

Differentially expressed genes (DEGs) were identified across all twelve clusters in reproductively old compared to young participants. The proportion of DEGs (adjusted p-value ≤0.0001) per genes tested ranged from 50.7% in granulosa cells to <5% in stromal and small lymphoid clusters (**Fig. 3f).** Notably the fold change of age-associated DEGs was highest in stromal cells compared to all other populations (mean |log2FC| = 2.5). Thus, stromal cells exhibit the most transcriptionally robust response associated with reproductive aging among all ovarian cell populations. Collectively, these data demonstrate that within the follicular fluid microenvironment, reproductive aging is accompanied by transcriptional remodeling that is disproportionately concentrated in stromal cells, occurring largely independent of shifts in cellular composition.

To further elucidate the cell types driving changes in ECM function and fibrosis, we analyzed a curated subset of fibroinflammatory genes across cell clusters. This gene list was derived from published transcriptomic and proteomic studies of ovarian aging and fibrotic disease in other organs, selecting genes with established roles in ECM remodeling, collagen regulation, cellular senescence, and inflammatory signaling.^18,19,28,61,63–68^ These genes were most differentially expressed in the stromal cluster with age relative to the macrophage, granulosa, and granulosa/theca clusters (**Fig 4a**). These results were further supported by unbiased analysis of the top 50 DEGs in stromal cells which revealed an age-associated upregulation of genes reflecting a fibrotic (*Bnc2, Fn1, Has1*) and contractile (*Myl9, Tpm1, Cldn1*) stromal phenotype alongside loss of epithelial cell adhesion programs (*Pcdh7, Ehf*; **Fig. 4b**). Gene Ontology (GO) pathway analysis of upregulated stromal DEGs identified enrichment of core matrisome genes, mesenchymal cell-cell adhesion, and ECM and tissue structure organization (**Fig. 4c**). Downregulated DEGs were enriched for pathways reflecting loss of cell identity (developmental cell lineages, epithelial cell differentiation, mucin O-glycosylation), mucosal innate immunity (acute-phase response, IL-17 signaling), and nuclear-receptor-mediated steroid hormone signaling, consistent with a shift away from tissue maintenance and homeostatic functions in the aged ovarian stroma (**Supplementary Fig. 3**). The fraction of ECM-related pathways was consistently enriched in stromal cells from reproductively old women across multiple analytical tools (**Fig. 4d**). Many of the genes driving these pathways that exhibited increased expression with age are key regulators of ECM fibrotic remodeling (*Col8a1*, *Serpine1*) and contraction (*Myh10*), pro-fibrotic fibroblast activation (*Igfbp5*, *Pdgfd*), and inflammation (*Il18*; **Fig. 4e**). Of note, the increased expression of *Col8a1* with age is consistent with the increase in PRO-C8 fragments observed in ECM-fingerprinting analysis (**Fig. 4e**; **Fig. 2d**). Collectively, these findings establish ovarian stromal cells as the primary drivers of fibroinflammatory transcriptional remodeling with reproductive aging, providing a molecular basis for the biomechanical stiffening detected by SWE in this cohort.

**Fig 4.**
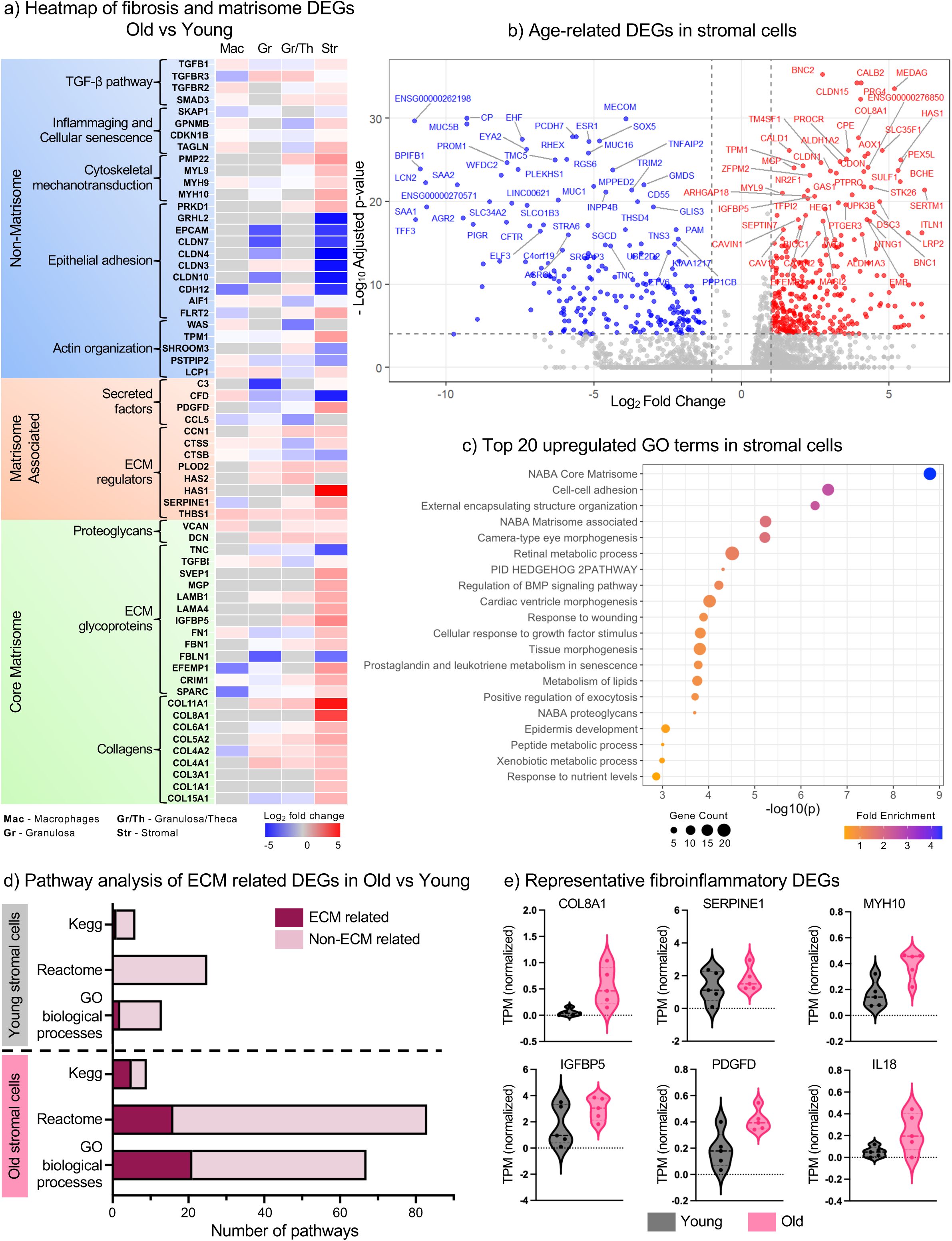
Aged ovarian stromal cells are enriched for fibroinflammatory and matrisome transcriptional programs. **a,** Heatmap of log₂ fold change of gene expression in reproductively old versus young participants for curated fibrosis, inflammaging, and ECM-regulatory genes across macrophage (Mac), granulosa (Gr), granulosa/theca (Gr/Th), and stromal clusters (Str) grouped by MatrisomeR category.^101^ **b,** Volcano plot of differentially expressed genes (DEGs) in old versus young participant stromal cells (adjusted p ≤0.0001). Colored points indicate genes meeting |log₂FC| ≥1.0 and adjusted p ≤0.0001 (Wilcoxon rank-sum with Bonferroni-correction) with top 50 genes labeled; red = upregulated in old (*n=* 363 DEGs), blue = downregulated in old (*n=* 225 DEGs), gray = not significant. **c,** Top 20 enriched GO biological process terms among upregulated stromal DEGs; bubble size represents gene count, color represents fold enrichment (low to high = yellow to blue). **d,** Number of ECM-related versus non-ECM-related pathways identified across KEGG, Reactome, and GO in young (top) versus old (bottom) stromal cells. **e,** Violin plots of normalized expression in transcripts per million (TPM) for representative fibroinflammatory DEGs in reproductively young and old stromal cells.

### Reproductive aging remodels intercellular communication toward a stromal-driven fibroinflammatory signaling network

Building on the identification of stromal cells as the primary drivers of age-related fibroinflammatory signatures, we applied CellChat to the scRNA-seq dataset to interrogate how reproductive aging reshapes intercellular communication within the ovarian microenvironment. First, we quantified the relative contribution of each cell type to the overall communication network, which revealed that stromal cells were the only cell type whose relative signaling contribution increased with age (**Fig. 5a**). Visualization of global communication networks centered on stromal-derived interactions highlighted that, while stromal cells engaged most cell populations in both age groups, interaction strength was markedly and selectively amplified toward immune clusters with age, indicating an enhanced stromal-to-immune signaling axis (**Fig. 5b**).

**Fig. 5.**
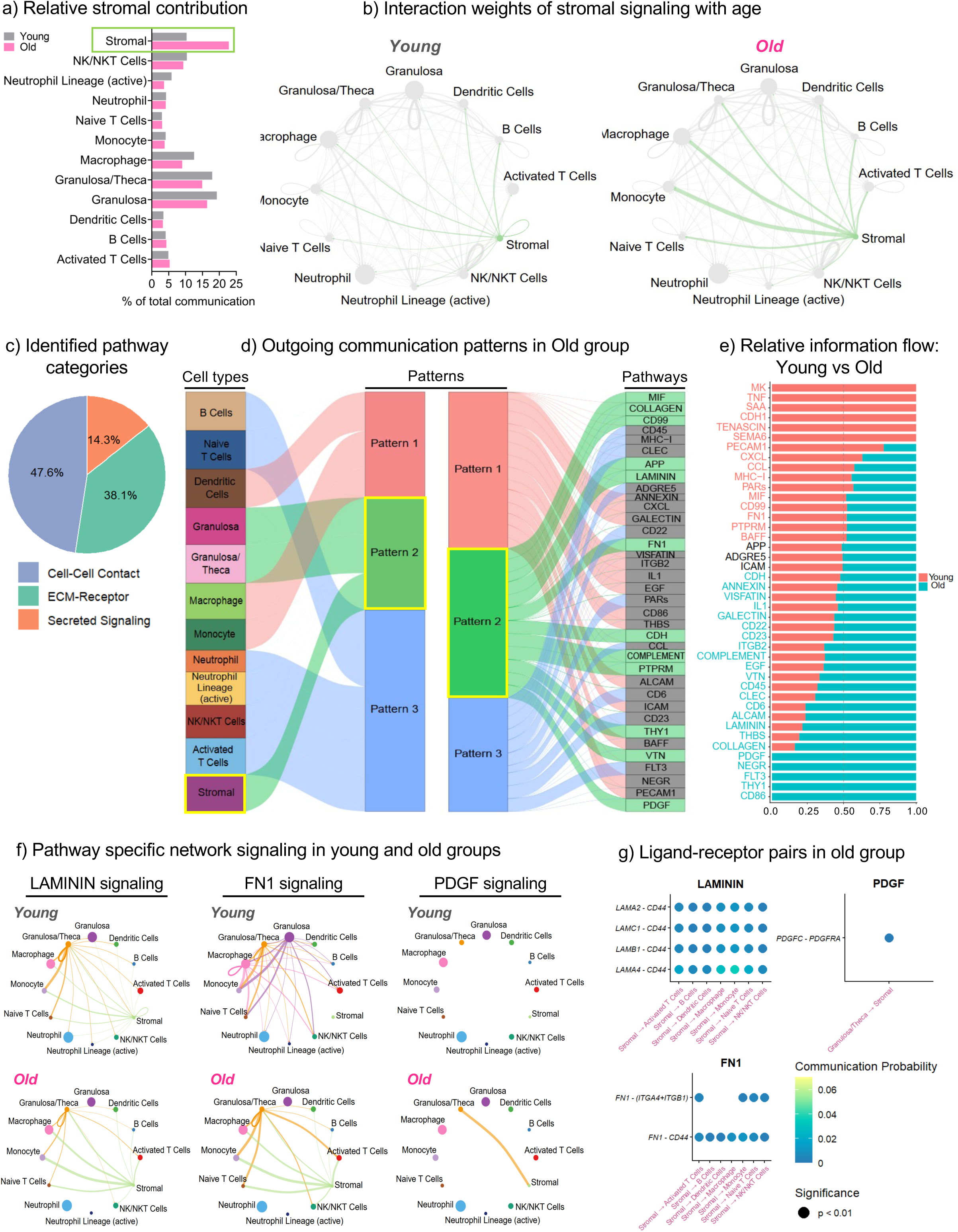
Aging enhances stromal-immune interactions and ECM-associated signaling in the ovarian microenvironment. **a,** The relative contribution of each cell type to the global cell-cell communication network in young and old groups. **b**, Global communication networks illustrating the strength of interactions between cell types in young and old populations. Edge thickness represents interaction weight; arrows indicate the direction of communication, highlighting the intensified stromal-to-immune axis in aging. **c,** Pie chart categorizing the identified signaling pathways into cell-cell contact, ECM-receptor, and secreted signaling modules. **d,** Heatmap showing the identification of three distinct signaling patterns. Pattern 2 represents the stromal-specific ECM-driven program dominant in the aged ovary. **e,** Comparison of the relative information flow (sum of communication probability) for significant signaling pathways between young and old groups. **f**, Pathway specific network visualization of *LAMININ*, *FN1* and *PDGF* signaling in reproductively young (top) and old (bottom) groups. **g,** Bubble plot identifying specific ligand-receptor pairs underlying selected pathways in the old group. Circle size represents the p-value; color intensity indicates the communication probability.

We classified the signaling pathways identified between cell types into three general categories: cell-cell contact, ECM-receptor interactions, and secreted signaling (**Fig. 5c**). ECM-associated pathways accounted for a large fraction of the total network, prompting us to look further into the specific signaling programs that drive this category. Analysis of outgoing signals in the reproductively old group revealed three distinct patterns across all identified cell types (**Fig. 5d**). Patterns 1 and 3 largely involved immune signaling pathways and immune cell populations (**Fig. 5d**). Pattern 2 was predominantly comprised of ECM and structural pathways and was associated with stromal cells, suggesting that the aged stroma tends to be more engaged sending out signals related to matrix-driven pathways (**Fig. 5d**). Consistent with this, comparison of the relative amount of signal coming from each age group among the top pathways revealed an aging-associated predominance of ECM-related pathways, including PDGF, COLLAGEN, and LAMININ (**Fig. 5e**). These data suggest that the aging stroma predominantly sends out signals related to ECM and these signals are largely unique to the reproductively old group.

The age-related shift towards ECM pathways was further validated by pathway specific network visualization of key ECM signaling pathways. The stroma was the major source of signaling for LAMININ and FN1 pathways which are ECM components that mediate cell adhesion and matrix organization. Although aggregate FN1 communication appeared comparable between age groups **(Fig. 5e),** this balance was driven by granulosa cell signaling in the young group. Cell-type-specific analysis revealed that stromal FN1 signaling was exclusive to the old group and completely absent in young stroma **(Fig. 5f).** The stroma also showed robust downstream LAMININ and FN1 signaling towards various immune cell types, including macrophages, monocytes, dendritic cells and lymphocytes (**left panel; Fig. 5f**). These stromal-borne immune interactions were significantly diminished or absent in the young group. In addition, ligand-receptor analysis demonstrated that the immune clusters predominantly expressed CD44 as the receptor for both stromal-derived LAMININ and FN1 (**Fig. 5g**). Given CD44-mediated binding toward ECM components is known to drive immune cell retention and inflammatory signaling^69^, our results suggest that stromal-immune interactions could play a role in chronic inflammation and matrix remodeling within aged ovarian microenvironment. In addition, PDGF signaling was detected specifically in the aged group with granulosa/theca cells signaling to stroma via PDGFRA (**Fig. 5f-g**). PDGF signaling activates MAPK and NF-κB pathways, while also inducing collagen production and matrix remodeling.^70^ Together, these data suggest a coordinated mechanism wherein ECM signaling retains immune cells within the stroma, while *PDGF* signaling drives fibrosis, establishing a pro-inflammatory and fibrotic ovarian microenvironment with reproductive age.

### Stiffness is a validated marker of ovarian aging independent of ovarian reserve

The corroboration of our biomechanical findings with molecular signatures suggests that ovarian stiffness reflects age-related fibroinflammatory remodeling. We thus sought to validate this relationship in a larger, prospectively enrolled cohort spanning a continuous age range and representing the broader clinical population seeking fertility care (Cohort 2, **Fig. 1a-i**). SWE was performed prospectively on 100 participants across reproductive age seeking fertility preservation or infertility treatment (**Fig. 1a-i**). Participants ranged from 24 to 45 years old (mean 35.1 ± 4.0) and data were collected for BMI, FSH, AMH, AFC, ovarian volume, ultrasound probe distance, and number of SWE measurements for each participant (**Supplementary Fig. 4a**). Participants in this study were predominantly non-Hispanic white women, reflecting the overall patient demographics in our fertility clinic (**Supplementary Fig. 4b**). Of the participants in the study, 57% were presumed fertile (seeking treatment for elective fertility preservation, male factor infertility, or another non-infertile reason) and 43% were infertile (uterine factor, diminished ovarian reserve, recurrent pregnancy loss, multiple factors, or unexplained infertility; **Supplementary Fig. 4c**). This cohort also exhibited positive correlations between age and FSH as well as negative correlations between age and both AMH and AFC (**Supplementary Fig. 4d-f**). These relationships are expected and demonstrate that our study subjects were broadly representative of the general population in terms of well characterized changes to the ovarian reserve. Presumed fertile and infertile individuals had overall similar clinical characteristics except for age being significantly higher (p = 0.03) in the infertile population, which reflects the expected impact of age on fertility (**Supplementary Table 2**).

To evaluate the relationship between age and ovarian stiffness parameters in this larger prospective cohort across a continuous spectrum of reproductive age, we performed a multivariable linear regression analysis. We analyzed the association of ovarian stiffness parameters with age when controlled for ovarian reserve (AMH and FSH), ovarian volume, BMI, and fertility status (**Table 3**). Age demonstrated a significant positive association with mean ovarian stiffness (β1 = 0.33; p = 0.03), elastic heterogeneity (β1 = 0.75; p = 0.01), and maximum stiffness (β1 = 0.93, p = 0.04, **Table 3**). Together, these multivariable analyses establish that the association between age and ovarian stiffness parameters persists independent of ovarian reserve, volume, BMI, and fertility status. Thus, mean ovarian stiffness, elastic heterogeneity, and maximum stiffness are distinct biomechanical indicators of ovarian aging. In addition, the strong association between ovarian stiffness and age persists in a larger, less tightly controlled cohort, suggesting broader applicability on a population-level and establishing fibrosis as a hallmark of human ovarian aging.

**Table 3.**
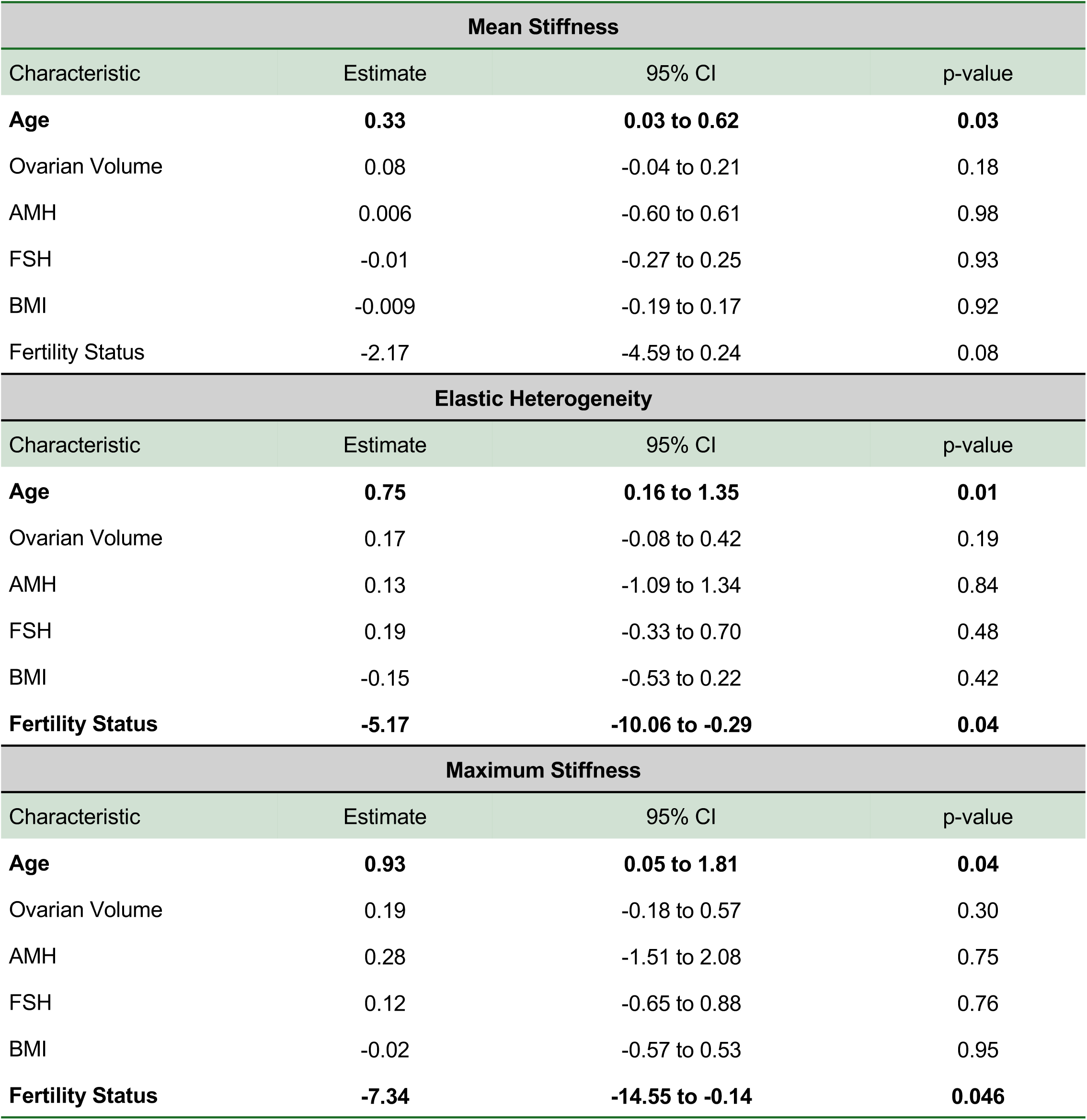
Association of ovarian stiffness with age in Cohort 2. Models were fit for the ovarian stiffness parameter (kPa) as a dependent variable, with age (years), ovarian volume (mL), AMH (ng/mL), FSH (ng/mL), BMI (kg/m²), and fertility status (presumed fertile versus infertile) as covariates. Estimates represent adjusted differences in stiffness (kPa).

## DISCUSSION

Ovarian fibrosis has emerged as a central feature of reproductive aging in animal models, but the human ovary has remained largely inaccessible to direct mechanistic interrogation, preventing the translation of compelling preclinical fibrosis biology into human reproductive medicine.^18–26,33,49,71,72^ By integrating transvaginal SWE, ECM neoepitope profiling, and single-cell transcriptomics within the same individuals, this study provides the first multimodal analysis of ovarian fibrosis in the human ovary in situ. We establish that human reproductive aging is accompanied by quantifiable ovarian stromal stiffening coupled to coordinated ECM remodeling, fibrotic stromal reprogramming, and increased stromal-immune crosstalk.

In our two-cohort design, age exerts an independent effect on ovarian stiffness, indicating that ovarian stiffness represents a stromal biomarker of the aging ovarian microenvironment not captured by conventional ovarian reserve markers that largely evaluate egg quantity, such as AMH or AFC. While BMI was modestly inversely associated with ovarian stiffness in Cohort 1, this relationship was not observed in the larger Cohort 2, suggesting that BMI is not a robust independent determinant of ovarian mechanical properties. Our observations strongly support prior animal and *ex vivo* studies that the ovary becomes biomechanically stiffer with age. Increased deposition of collagen matrices and hyaluronan depletion in aged mouse and fixed human ovary stroma have been directly associated with a quantifiably stiffer tissue.^18,20,22^ EH emerged as one of the strongest age-associated ovarian stiffness parameters across both cohorts in our study, suggesting that focal spatial variability in stromal architecture is a key biomechanical feature of ovarian aging. This observation aligns closely with prior histologic studies describing collagen-rich fibrotic foci throughout the aging ovarian stroma rather than uniform matrix deposition.^18,20,23,25^ Notably, increased ovarian stiffness independent of age was also associated with diminished ovarian responsiveness to hormonal stimulation during assisted reproduction, reduced numbers of follicles maturing to the ovulatory stage (lower FORT), and lower oocyte yield from antral follicles (lower FOI). These data corroborate *in vivo* animal and *in vitro* follicle growth studies where increased matrix stiffness impairs folliculogenesis and leads to dysfunctional ovulation with increased retained oocytes.^22,29^ Although additional outcome data such as euploidy incidence and live birth rates are needed, these findings support stromal stiffness as a functional determinant of ovarian competence rather than simply an epiphenomenon of aging.

Our study provides the first application of ECM neoepitope profiling to reproductive aging. Whereas prior ovarian fibrosis studies have relied predominantly on static histologic measurements, ECM neoepitope analysis provides dynamic insights into fibroproliferation and revealed evidence of disrupted matrix homeostasis with aging. Across paired serum and follicular fluid, reproductive aging was characterized by a coordinated shift toward matrix accumulation primarily attributable to collagen VIII formation and reduced collagen VI degradation. Among all analytes, PRO-C8 demonstrated the most pronounced age-associated enrichment, suggesting collagen VIII remodeling as a central feature of ovarian fibroinflammaging. Collagen VIII is recognized as an injury-associated matrix component implicated in vascular remodeling, fibrosis, and tissue stiffening in cardiovascular, renal, and pulmonary disease.^54,73–78^ Unlike fibrillar collagens that primarily provide structural bulk to the ECM, collagen VIII is stress-responsive and participates in matrix organization, endothelial stability, fibroblast activation, and TGF-β– associated remodeling^73,74,76^ PRO-C8 targets the vastatin domain, which is the C-terminal NC1 fragment of the collagen VIII α1 chain. Serum vastatin correlates significantly with age in healthy individuals of mixed sex.^79^ The parallel upregulation of stromal *Col8a1* expression in reproductively older ovaries provides transcriptomic validation of our proteomic findings and identifies collagen VIII as a candidate mediator of age-associated ovarian fibrosis. In contrast, lower follicular fluid C6M with reproductive age further suggests reduced collagen VI turnover and progressive matrix accumulation. Dysregulated collagen VI remodeling has emerged as a central feature of organ fibrosis^80,81^ and circulating endotrophin (PRO-C6) independently predicts a 2-fold increase in all-cause mortality.^38^ Together, these findings suggest that ovarian aging is characterized not simply by collagen accumulation, but by broader disruption of ECM homeostasis favoring matrix persistence and tissue stiffening. The intraindividual concordance between serum and follicular fluid neoepitope profiles suggests that circulating ECM biomarkers may capture biologically meaningful remodeling within the ovarian microenvironment. If replicated in larger cohort studies, this highlights ovarian fibrosis could ultimately be monitored using minimally invasive circulating ECM neoepitope biomarkers, analogous to emerging fibrosis frameworks in hepatic^36,82^, renal^83,84^, and cardiovascular pathologic aging.^36^

Single-cell transcriptomics of follicular fluid aspirates revealed that stromal cells are the dominant drivers of ovarian fibroinflammaging. The highest upregulated genes in the stroma of reproductively old participants have established roles in fibrosis, cellular senescence, and chronic inflammatory remodeling across aging tissue and fibrotic diseases. *Fn1* serves as a foundational scaffold for fibrotic ECM assembly, that enhances recruitment of latent TGF-β-binding protein-1 to the fibroblast matrix^85^ and drives profibrotic fibroblast activation via integrin α5β1-mediated mechanotransduction.^86,87^ *Serpine1* mediates TGF-β1-induced cellular senescence and SASP-driven macrophage activation^88^ while *Igfbp5* propagates senescence through paracrine signaling and ROS production.^89^ Lastly, *Pdgfd* drives fibroblast proliferation and collagen secretion via TGF-β1 positive feedback in numerous fibrotic diseases^70,90,91^, with its knockout attenuating fibrosis in injury models.^92^ Aging amplified stromal-to-immune communication and shifted signaling architecture toward ECM-dominant pathways including FN1, LAMININ, COLLAGEN, and PDGF signaling, consistent with proteomic analyses demonstrating coordinated upregulation of ECM remodeling and immune response pathways in the aging mouse ovary^19^. The prominence of CD44-mediated stromal–immune interactions is particularly compelling as CD44 mediates TGF-β activation through MMP-dependent mechanisms, promotes fibroblast migration, immune retention, chronic inflammation, and drives myeloid-derived suppressor cell induction.^69,93,94^ The observation that fibroinflammatory pathways predominate despite the overall small sample size of patients used for single-cell sequencing suggests that this finding is robust. These senescence- and fibrosis-associated programs within the aged ovarian stroma suggest that the ovary undergoes fibroinflammatory remodeling similar to fibrosis in other organs and lays the foundation for therapeutic targeting.

Collectively, our data position ovarian fibrosis not as a byproduct of reproductive aging, but as a central and measurable biological process linked to deteriorating ovarian function. These findings have substantial translational implications. Anti-fibrotic strategies have recently emerged as promising interventions for ovarian aging that improve follicular function in preclinical models through collagen depletion approaches^24^, metformin-mediated immune mediation^25^, modulating inflammatory pathways^49^, targeting metabolic homeostasis^95^, and systemic anti-fibrotic therapies.^30,96^ However, real-world translation has been constrained by the inability to demonstrate that age-related ovarian stiffening is a robust phenotype in humans due to the absence of non-invasive tools capable of monitoring human ovarian fibrosis longitudinally *in vivo*. Our findings conclusively demonstrate this phenotype in humans for the first time and support transvaginal SWE as a novel and clinically tractable platform to fill this gap. By enabling direct assessment of ovarian biomechanical properties across the reproductive lifespan, SWE may provide both a mechanistic biomarker of ovarian aging and a quantitative endpoint for future anti-fibrotic clinical trials in women for ovarian aging. Future longitudinal studies may continue to identify additional value of ovarian stiffness measurements, including possible relationships with live birth outcomes or menopause timing.

In summary, this study is the first to integrate biomechanical, paired serum and follicular fluid proteomic, and transcriptomic profiling within the same individuals to provide unique characterization of human ovarian fibroinflammaging. This study establishes ovarian fibrosis as a reproducible, measurable hallmark of human reproductive aging and identifies stromal fibroinflammatory remodeling as a central biological feature of ovarian decline. Through multimodal integration of biomechanics, ECM fingerprinting, and single-cell transcriptomics, we define a translational framework for minimally invasive detection of ovarian aging and provide mechanistic support for targeting fibrosis as a potentially modifiable determinant of human reproductive longevity.

## METHODS

### Study design and ethical approval

This study employed a prospective, multimodal design to characterize biomechanical, molecular, and transcriptomic features of human ovarian aging across two independent cohorts enrolled at a single academic fertility center, Northwestern Medicine Center for Fertility and Reproductive Medicine **(Fig. 1a).** All participants provided written informed consent. Study protocols were approved by the institutional review board (IRB); Cohort 1 (STU00222297) and Cohort 2 (STU00218882).

### Cohort 1 – Age stratified cohort

Cohort 1 comprised of participants undergoing planned fertility preservation or in vitro fertilization (IVF), stratified into two discrete reproductive age groups: Reproductively Young (age ≤33 years) and Reproductively Old (age ≥37 years) **(Fig. 1a).** Cohort 1 participants were recruited based on strict criteria designed to isolate the effects of ovarian aging while minimizing potential confounding from systemic and ovarian pathologies. The reproductively young group (age ≤33 years) was restricted to individuals undergoing planned fertility preservation or IVF treatment due to male-factor infertility or a same-sex female relationship. The reproductively old group (age ≥37 years) also included infertility diagnoses of diminished ovarian reserve (DOR) or unexplained infertility. In the latter “unexplained” subgroup, a comprehensive evaluation, including assessment of ovarian reserve, tubal patency, uterine cavity, and semen parameters, was normal, thereby supporting ovarian aging as the primary underlying etiology. We excluded diagnoses that may independently impact the ovary including ovulatory dysfunction, polycystic ovarian syndrome (PCOS), endometriosis, diabetes mellitus, active or prior malignancy, or autoimmune disease. Body mass index (BMI) was restricted to 19 to 30 kg/m², and participants with a history of ovarian surgery, tobacco or substance use, or a prior oocyte retrieval within the preceding 12 months were also ineligible. Given the potential for exogenous hormones to alter the ovarian microenvironment, we also excluded participants on long-term systemic hormonal contraception or ovarian stimulation protocols requiring estrogen priming or oral contraceptive pretreatment for more than 21 days. All study participants underwent ovarian stimulation with either GnRH antagonist or microdose GnRH agonist flare protocol.

All participants underwent early follicular phase (cycle day 2–3) transvaginal ultrasound with SWE and concurrent serum collection at the start of ovarian stimulation **(Fig. 1b).** Antral follicle count (AFC), total ovarian volume, ultrasound probe distance (average distance of probe- to- ovary) and baseline hormonal labs were obtained on the same day as SWE. Most recent anti-mullerian hormone (AMH) and follicle stimulating hormone (FSH) were also recorded. At the time of oocyte retrieval, otherwise discarded follicular aspirate was collected from three dominant follicles per ovary. A subset of Cohort 1 participants underwent ECM fingerprinting of follicular fluid and matched serum, as well as single-cell RNA sequencing (scRNA-seq) of follicular fluid **(Fig 1c).**

Participant demographic data and ART cycle details including total gonadotropin dose were recorded. The primary stimulation response outcomes were total number of oocytes retrieved, total number of mature (metaphase II; MII) oocytes, and maturation rate (MII/total oocytes). Two composite ART performance indices were calculated: 1) the follicular output rate (FORT), defined as the ratio of follicles measuring ≥ 15mm on day of trigger to the total AFC at start of stimulation, and 2) the follicle-to-oocyte index (FOI), defined as the ratio of total oocytes retrieved to the total AFC.^97,98^

### Cohort 2 – Continuous age-distributed cohort

Cohort 2 was designed to validate and extend the biomechanical findings from Cohort 1 by examining the relationship between continuous chronological age and ovarian stiffness in a larger, real-world clinical population across the reproductive lifespan. This cohort included individuals aged 24 to 45 years presenting for diagnostic evaluation prior to fertility treatment over an 18-month period **(Fig. 1a).** Participants were excluded if they had undergone oocyte retrieval within the preceding 12 months or had conditions known to independently alter ovarian stiffness, including dermoid cysts, endometriomas, or PCOS. All participants underwent early follicular phase (cycle days 2–3) transvaginal ultrasound with SWE during their initial diagnostic evaluation. Ultrasound probe-to-ovary distance measurement and AFC was also collected at the same time. Participants were included in the analysis if at least three SWE measurements were obtained within high-confidence regions of a single ovary. Cohort 2 participants were distinct from those enrolled in Cohort 1.

Demographic and clinical data was also collected for each participant, including age, BMI, AMH and FSH levels, antral follicle count (AFC), ovarian volume, fertility diagnosis, race, and ethnicity. Participants were categorized as “presumed fertile” or “infertile” based on their clinical history and reasons for seeking fertility counseling. Presumed fertile patients were seeking fertility counseling either for planned fertility preservation, male infertility, or another non-infertile reason (such as same-sex couples). Patients seeking treatment for infertility were categorized based on the Society for Assisted Reproductive Technologies (SART) criteria.^99^

### Transvaginal 2D Ultrasound with Shear Wave Elastography

Participants underwent transvaginal ultrasonography using a GE Logic Fortis HDU ultrasound device (SN LFO301952, GE HealthCare) with a 2D transvaginal probe (IC5-9-D; SN 1278929WX3, frequency range: 7-9 MHz) **(Fig.1b).** For Cohort 1, all ultrasound examinations were performed by a single sonographer to ensure measurement consistency at the baseline visit for ovarian stimulation. In the larger Cohort 2, scans were conducted by a team of five sonographers with expertise in gynecologic ultrasonography at time of diagnostic ultrasound during initial fertility evaluation at our clinic. Least possible compression was applied intravaginally such that the ultrasound probe was within 2 cm of depth of the ovary. Up to twelve SWE frames in a 5 mm region of interest (ROI) were acquired in high-confidence regions in each ovary, as determined by a color-coded confidence map **(Supplementary Fig. 1).** High confidence regions typically consist of areas devoid of fluid-filled follicles and any other ovarian pathology. The procedure was completed for both ovaries whenever possible. Any measurements taken in ROIs that included low-confidence areas were excluded from analysis. Multiple stiffness parameters were extracted for analysis. For each individual, all measurements taken across both ovaries were grouped together to calculate the average stiffness (reported as mean stiffness) and to identify the single measurement with the highest stiffness value (reported as maximum stiffness). We also quantified heterogeneity of the measurements in each participant using elastic heterogeneity as a measurement, which is calculated as the average of the highest three stiffness values minus the average of the lowest three stiffness values **(Fig. 1b)**.^38^ Elastic heterogeneity has been previously reported to improve diagnostic performance of SWE for identification of malignant breast lesions.^38^

### Follicular Fluid and serum collection

For Cohort 1, follicular fluid and serum were collected from 20 participants per age group for ECM fingerprinting. In a subset, follicular fluid from 5 participants per age group was additionally processed for single cell RNA sequencing (scRNAseq) **(Fig. 1c**). Controlled ovarian stimulation was performed using either a GnRH antagonist or microdose GnRH agonist flare protocol at the discretion of the treating physician. Final oocyte maturation was triggered with GnRH agonist, human chorionic gonadotropin (hCG), or a combination of both. Follicular fluid and cellular aspirates were collected from three dominant follicles (≥18mm) per ovary at the time of transvaginal oocyte retrieval. Samples were centrifuged at 500 × g for 10 minutes at 4°C. Cell pellets were isolated for scRNA-seq processing, and protease inhibitor (1:100 from 100× stock; Thermo Fisher Cat. No. 87786) was added to the supernatants immediately after centrifugation before aliquoting and storage at −80°C for ECM neoepitope profiling. Matched serum samples were collected on the same morning as the SWE visit and processed within 2 hours of collection. Serum was centrifuged at 1,000 × g for 15 minutes, aliquoted, and stored at −80°C.

### Extracellular matrix fingerprinting

To quantify ECM remodeling associated with fibrosis and fibrolysis, a panel of circulating and follicular fluid ECM neoepitope biomarkers was measured in 20 participants per age group from Cohort 1. Follicular fluid from the right and left ovaries was pooled prior to analysis. In total, 11 biomarkers of ECM degradation, and ECM formation were measured **(Fig 2a).** Seven markers related to ECM degradation: type I, III, and IV, VI collagen degradation (nordicC1M^TM^, nordicC3M^TM^, nordicC4M^TM^, nordicCAN^TM^, nordicC6M^TM^, respectively), MMP-degraded mimecan (nordicMIM^TM^), and MMP-degraded biglycan (nordicBGM^TM^). Four markers related to ECM formation: type I, III, type VI and VIII collagen (nordicPRO-C1^TM^, nordicPRO-C3^TM^, nordicPRO-C6^TM^, and nordicPRO-C8^TM^, respectively). All biomarkers were evaluated by ELISA measurements and quantified according to the manufacturer’s instructions (Nordic Bioscience A/S, Herlev, Denmark; **Supplementary Table 3**). Each ELISA plate included samples from each of the different age groups in duplicates. Samples were re-assayed if the coefficient of variation (CV) between the duplicates was higher than 15%.

Log₂ fold-change heatmaps were generated as log₂(mean Old / mean Young) to visualize the magnitude and direction of age-associated differences of each neoepitope across compartments. Intra-individual correlations between serum and follicular fluid levels were assessed for each biomarker.

### Single cell isolation from follicular aspirates

Single-cell suspensions were generated from fresh follicular fluid as previously described by our group.^100^ Briefly, pooled right and left ovary follicular aspirates were processed within 2–3 hours of oocyte retrieval and subjected to sequential centrifugation and wash steps. Cell pellets were enzymatically dissociated using dispase and DNase I, followed by trypsin digestion, and filtered through 100 μm and 40 μm strainers to obtain a single-cell suspension. Red blood cells were removed using ACK lysis buffer, and cells were washed and resuspended in PBS-based buffer. Cell number and viability were assessed prior to sequencing; all samples had viability >70% and cell concentrations within the optimal range (1×10⁵ to 2×10⁶ cells/mL).

### Single cell RNA sequencing

Cell number and viability were analyzed using Nexcelom Cellometer Auto2000 with AOPI fluorescent staining method. Sixteen thousand cells were loaded into the Chromium X Controller (10X Genomics, PN-1000328) on a Chromium GEM-X Single cell 3’ Chip Kit v4 (10x Genomics, PN-1000215) and processed to generate single cell gel beads in the emulsion (GEM) according to the manufacturer’s protocol. The cDNA and library were generated using the GEM-X Single cell 3’ Kit v4 (10x Genomics, PN-1000691) and Dual Index Kit TT Set A (10X Genomics, PN-1000215) according to the manufacturer’s manual. Quality control for constructed library was performed by Agilent Bioanalyzer High Sensitivity DNA kit (Agilent Technologies, 5067-4626) and Qubit DNA HS assay kit for qualitative and quantitative analysis, respectively. The multiplexed libraries were pooled and sequenced on Illumina Novaseq X Plus sequencer with 100 cycle kits using the following read length: 28 bp Read1 for cell barcode and UMI, and 90 bp Read2 for transcript.

### Single cell RNA sequencing and analysis

Raw sequencing data, in base call format (.bcl) was demultiplexed using Cell Ranger from 10x Genomics, converting the raw data into FASTQ format. Cell Ranger was also used for alignment of the FASTQ files to the human reference genome (hg38) and count the number of reads from each cell that align to each gene. The resulting matrix files which summarize the alignment results were imported in Seurat (Satija Lab, NYGC) for further analysis. In Seurat, each individual sample was preprocessed, normalized, and scaled using SCTransform. Each sample was checked with quality control measures for the number of genes, counts, and percent mitochondrial genes detected per cell, and these results were used to determine thresholds to remove outlying cells. All samples were combined into a single dataset, adding appropriate metadata to each sample. After integrating the samples, the cells were clustered using the Seurat tools FindNeighbors, FindClusters, and RunUMAP using default parameters with the exception of dims=12. Based on the results of FindClusters, we settled on a resolution of 0.2 for the rest of the analysis.

### Cell type identification

To assist in cell type determination, we used the Seurat FindAllMarkers function based on a Wilcox likelihood-ratio test with default parameters, and selected the genes expressed in more than 25% of the cells in a cluster with an average log(fold change) value greater than 0.25 as DEGs. For the cell type annotation of each cluster, we combined the expression of canonical markers found in the DEGs between clusters with knowledge from the literature and displayed the expression of markers for each cell type using a dot plot generated in R. We next compared the average cell proportion per each cluster to the total average population in the reproductively young (n=5) and reproductively old participants (n=5).

### Reproductive old versus young DEG and enrichment pathway analysis

Differential gene expression analyses were performed between reproductively old and young samples, separately for cell type, using Seurat FindMakers with only.pos=FALSE, logfc.threshold = 0, thresh.use = 0 and the default min.pct = 0.01. DEGs with adjusted p-value ≤ 0.0001 were classified into core matrisome (collagens, ECM glycoproteins, proteoglycans) and matrisome-associated (ECM regulators, ECM-affiliated proteins, secreted factors) categories using the *Homo sapiens* matrisome annotation in Matrisome AnalyzeR.^66,101^ Additional categories of recognized genes involved in inflammaging and cellular senescence, and the TGF-β pathway were also identified.^63,102^ Gene Ontology (GO), Reactome, and KEGG analyses were performed to explore unbiased functional enrichment of the significant age related DEGs with adjusted p-value ≤0.0001.^103–106^ All pathway-related genes mentioned in this manuscript were obtained by the combination of related terms from the GO databases.^103–106^ Heatmaps were generated using GraphPad Prism 8.0. Volcano plots were generated in R. For visualization purposes, volcano plots highlighted genes with an adjusted p-value ≤ 0.0001 and |log₂FC| ≥ 1. Violin plots of representative fibroinflammatory DEGs were created using GraphPad Prism 8.0.

### Cell-cell communication analysis

Cell-cell communication analysis was performed using the CellChat R package (v.1.6.1) on the scRNA-seq dataset^107^. Separate CellChat objects were generated for young and old groups from the Seurat object. Communication probabilities were inferred based on known ligand-receptor interactions and average gene expression within each cell type. The inferred communication networks were used to compare overall interaction strength, the contribution of different cell types, and signaling pathway activity between young and old groups. Signaling pathways were further analyzed by comparing relative information flow and identifying outgoing communication patterns. Selected pathways, including LAMININ, FN1, and PDGF, were examined at both the pathway and ligand-receptor levels. Cell interactions with a p-value < 0.05 were considered significant. All visualizations of cell-cell communication networks were generated using functions implemented in CellChat.

### Statistical analysis

All statistical analyses were completed using GraphPad Prism 10 and in consultation with the Northwestern Biostatistics Collaboration Center. Measurements following a normal distribution were reported as mean ± standard deviation. Unpaired T-test and one-way ANOVA were used to make comparisons between groups. Pearson’s correlation analysis was completed between two variables and is reported as a correlation coefficient (r) and p-value. Multivariable linear regressions were completed with stiffness parameters (mean stiffness, elastic heterogeneity, or maximum stiffness) as dependent variables and age, ovarian volume, AMH levels, FSH levels, BMI, and fertility status (categorized as presumed fertile or infertile) as independent variables. AFC was excluded from the multivariable linear regression analysis due to high levels of multicollinearity with AMH. Results were reported as the regression estimate, 95% confidence interval, and p-value for each variable. For all statistical analysis, significance was set at P < 0.05.

## ACKNOWLEDGEMENTS

We thank Cecilie Møller Hausgaard for performing the extracellular matrix fingerprinting assays on the serum and follicular fluid samples included in this study.

---

**Supplementary Fig. 1.**
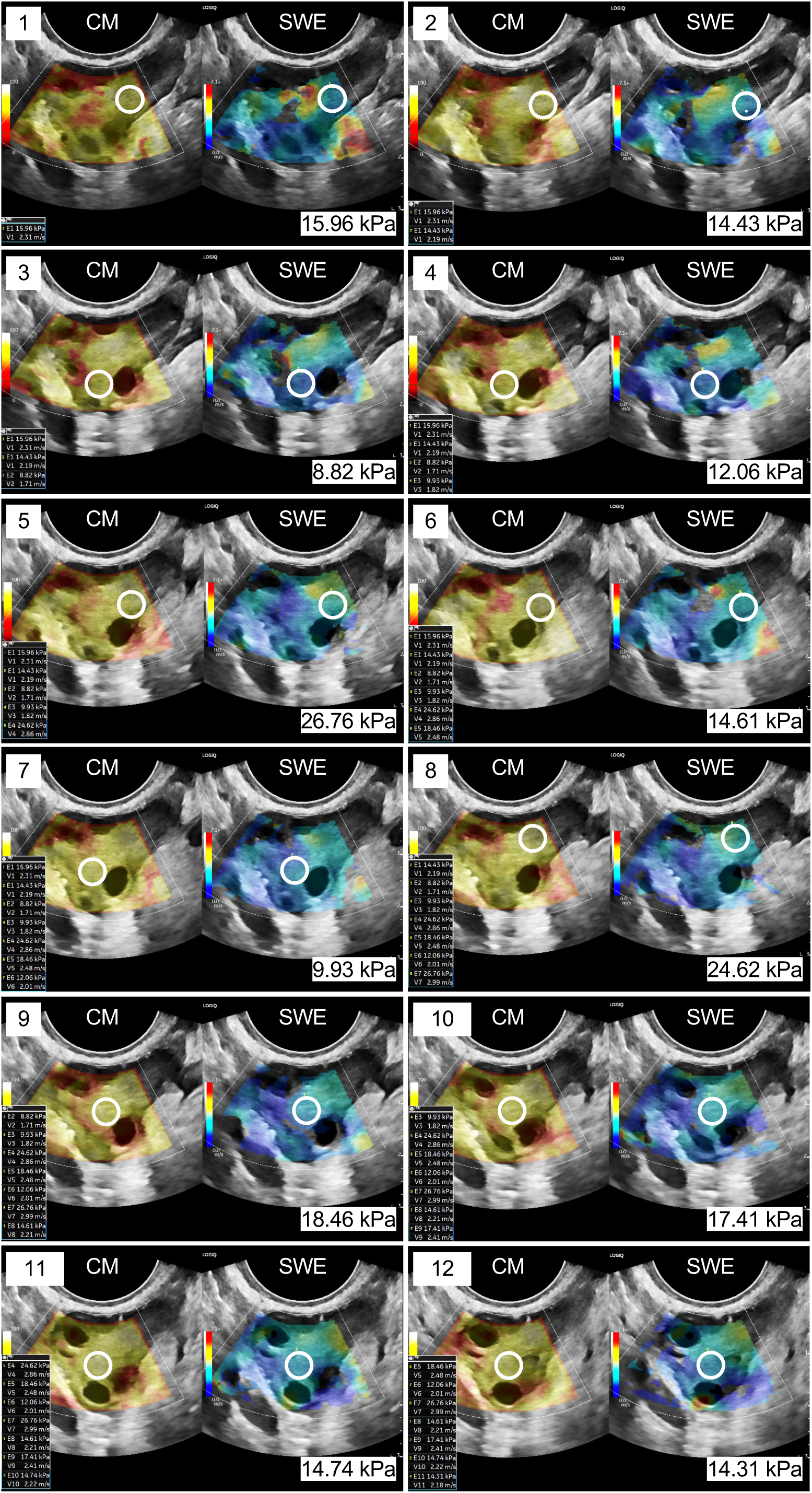
Representative images and stiffness measurements from one ovary of an individual participant from Cohort 1. The image on the left for each measurement represents the confidence map (CM) with yellow and white regions indicating high confidence areas. The image on the right for each measurement represents the shear wave elastography (SWE) stiffness measurement. The white circle represents the 5mm region of interest (ROI) corresponding to each stiffness (kPa) measurement.

**Supplementary Fig. 2.**
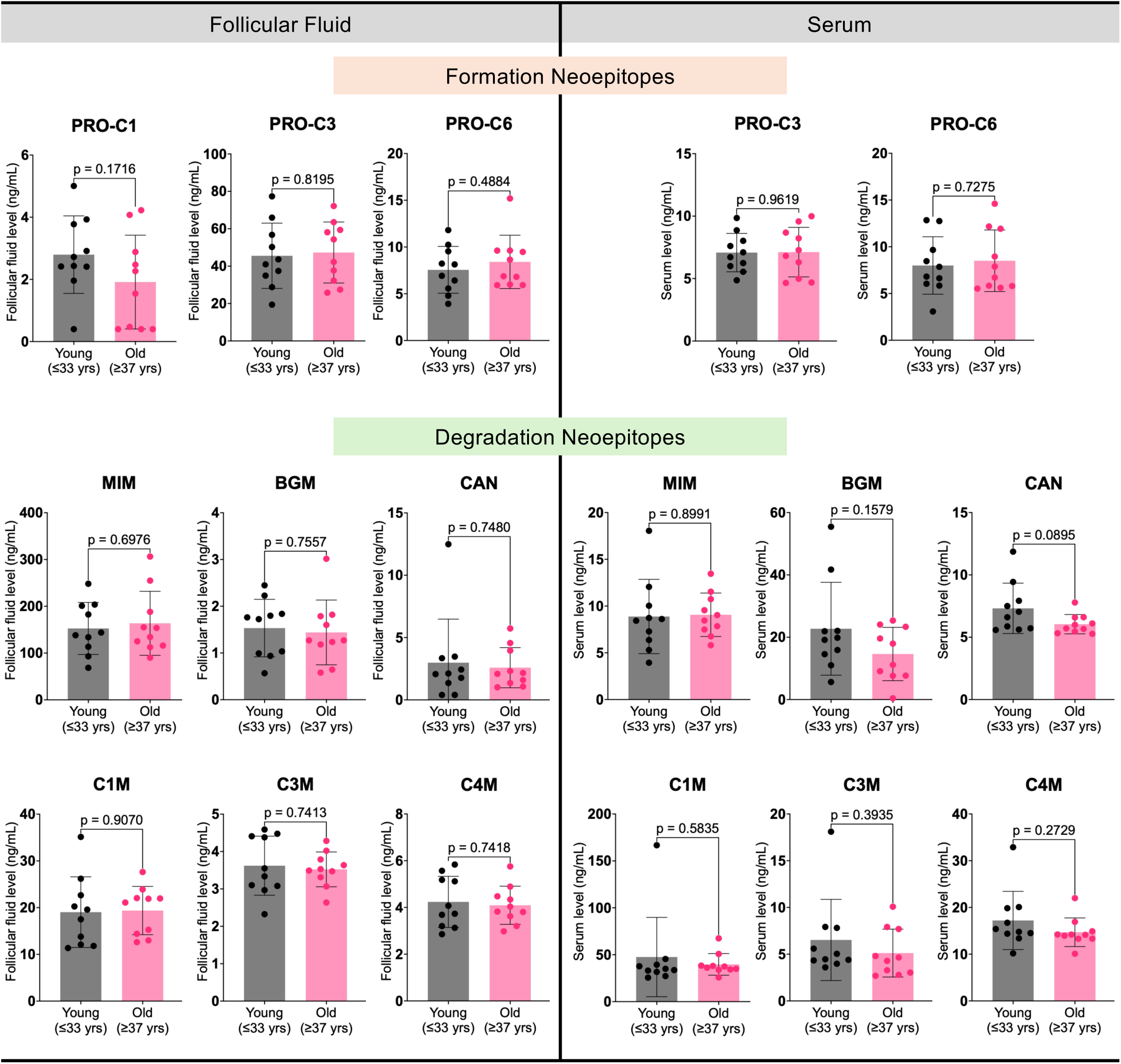
Additional ECM neoepitopes in follicular fluid and serum by age group. Bar graphs of additional formation (top) and degradation (bottom) ECM fingerprinting neoepitopes analyzed in reproductively young (grey; *n=* 10) compared to old participants (pink; *n=* 10). Paired follicular fluid (left) and serum (right) was collected from each participant. Each dot represents a participant. Bars represent mean ± SD. Labeled p-values by unpaired t-test.

**Supplementary Fig. 3.**
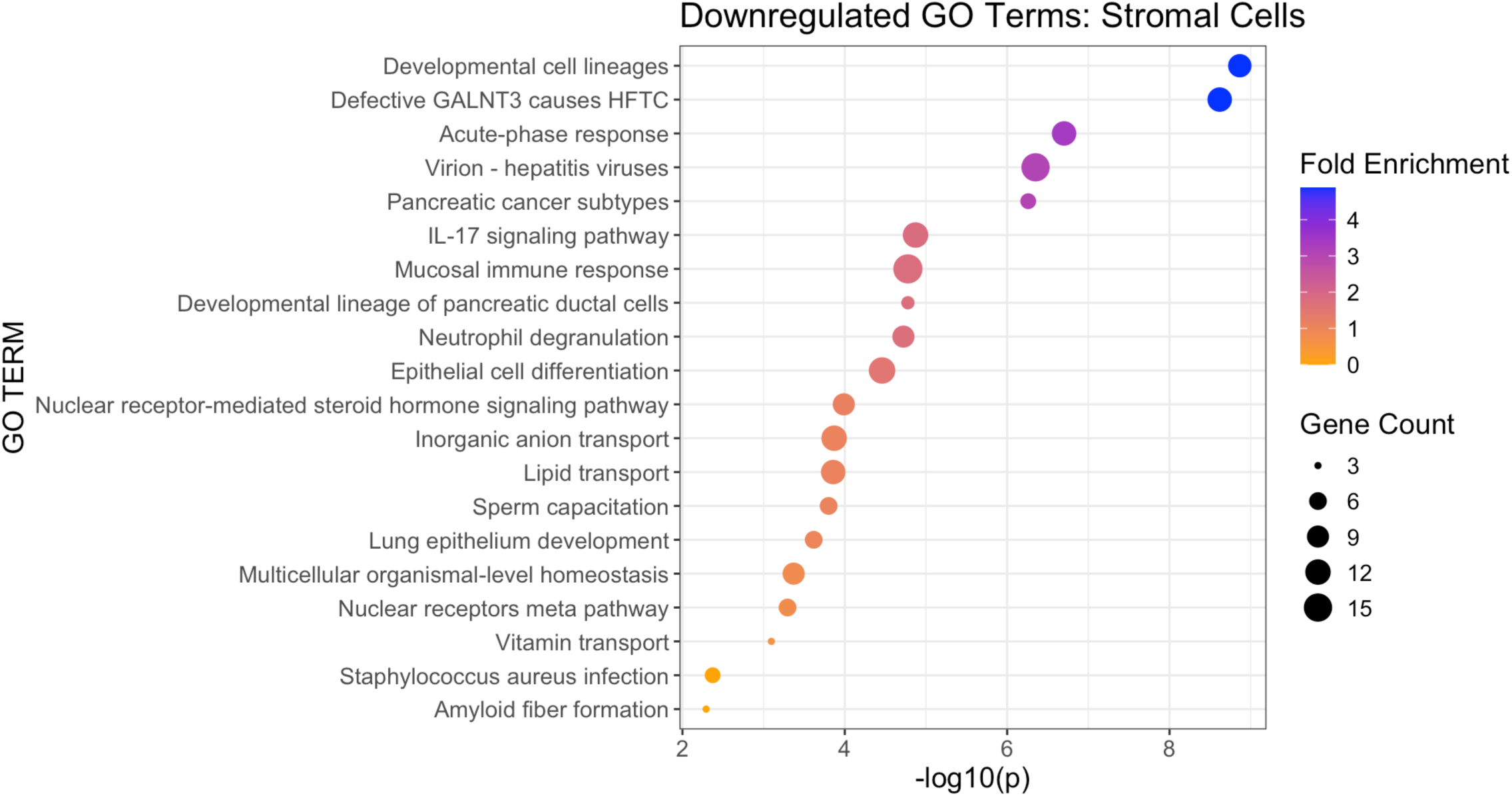
Stromal cell downregulated GO terms with reproductive age. Top 20 GO biological process terms among downregulated stromal differentially expressed genes (DEGs) in single cell transcriptomic profiling of ovarian follicular fluid aspirates from reproductively old (*n=*5) compared young (*n=*5) participants. Stromal DEGs with adjusted p≤0.0001. Bubble size represents gene count. Color represents fold enrichment (low to high = yellow to blue).

**Supplementary Fig. 4.**
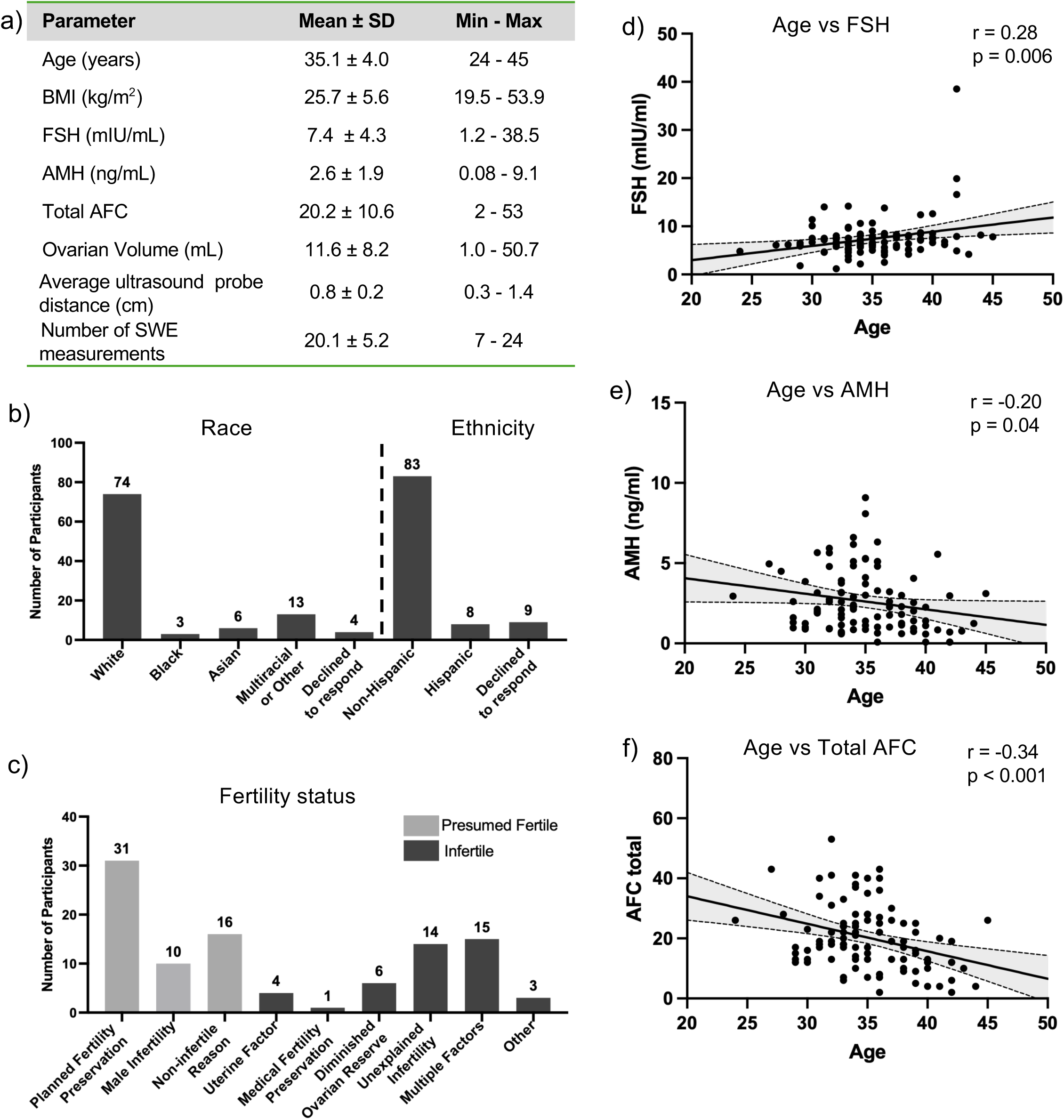
Cohort 2 demographic information. **a,** Cohort 2 of continuous age-distributed participant demographic information *(n=* 100; 24-45 years old). **b,** Participants self-reported racial identity and ethnicity. **c,** Overview of participant indications for seeking fertility counseling. **d,** Participants exhibit expected correlations with age and FSH (positive; r= 0.30, p= 0.002), **e,** AMH (negative; r= −0.29, p= 0.004), and **f,** AFC (negative; r= −0.36, p <0.001).

**Supplementary Table 1.** Cohort 1 demographic information. Reproductive young versus old group analysis by unpaired t-test.

| Characteristic | Reproductively Young<br>≤33 years (n=16) | Reproductively Old<br>≥37 years (n=16) | p-value |
| --- | --- | --- | --- |
| **Age (years)** | 30.5 ± 1.7 | 40.3 ± 2.0 | **<0.000001** |
| **BMI (kg/m<sup>2</sup>)** | 22.5 ± 1.9 | 24.5 ± 2.6 | **0.02** |
| FSH (mIU/mL) | 6.2 ± 2.4 | 7.5 ± 2.9 | 0.18 |
| **AMH (ng/mL)** | 3.1 ± 1.7 | 1.7 ± 1.1 | **0.01** |
| **Total AFC** | 24.4 ± 8.4 | 12.8 ± 5.1 | **<0.0001** |
| **Total ovarian volume (mL)** | 13.9 ± 4.8 | 10.1 ± 4.9 | **0.03** |
| Average US probe distance (cm) | 0.7 ± 0.4 | 0.8 ± 0.2 | 0.68 |
| Number of SWE measurements | 24 ± 0 | 23.2 ± 3 | 0.33 |
| Total FSH Dose (IU) | 3861 ± 1312 | 4659 ± 1206 | 0.08 |
| **Total retrieved oocytes** | 20 ± 12 | 11 ± 7 | **0.02** |
| **Total mature (MII) oocytes** | 15 ± 10 | 8.6 ± 5.6 | **0.03** |
| Percent MII oocytes (%) | 74.1 ± 17 | 76.0 ± 17 | 0.75 |
| Cycle Diagnosis (n=) |  |  |  |
| *Planned fertility preservation* | 9 | 2 |  |
| *Male infertility* | 6 | 2 |  |
| *Diminished ovarian reserve* | - | 7 |  |
| *Unexplained infertility* | - | 4 |  |
| *Non-infertile indication* | 1 | 1 |  |

**Supplementary Table 2:** Custom panel of ECM neoepitope biomarkers.

| Biomarker | Description of measure | Reference |
| --- | --- | --- |
| Tissue Degradation |  |  |
| C1M | MMP-2, -9, -13 mediated degradation of type I collagen | (D. Leeming et al., 2011) |
| C3M | MMP-9 mediated degradation of type III collagen | (Barascuk et al., 2010) |
| C4M | MMP-2, -9, -12 mediated degradation of type IV collagen | (Sand et al., 2013) |
| CAN | Cathepsin-S mediated degradation of type IV collagen | (S. H. Nielsen et al., 2025) |
| C6M | MMP-2, -9 mediated degradation of type VI collagen | (Veidal et al., 2011) |
| MIM | MMP-9 -degraded mimecan | (Barascuk et al., 2011) |
| BGM | MMP-9 degraded biglycan | (Genovese et al., 2013) |
| Tissue Formation |  |  |
| PRO-C1 | Pro-peptide of type I collagen | (D. J. Leeming et al., 2010) |
| PRO-C3 | N-terminal propeptide of type III collagen | (M. J. Nielsen et al., 2013) |
| PRO-C6 | C-terminal NC domain of type VIa3 collagen (Endotrophin) | (Sun et al., 2015) |
| PRO-C8 | C-terminal domain of type VIII collagen (Vastatin) | (Ulrik et al., 2016) |

**Supplementary Table 3.**
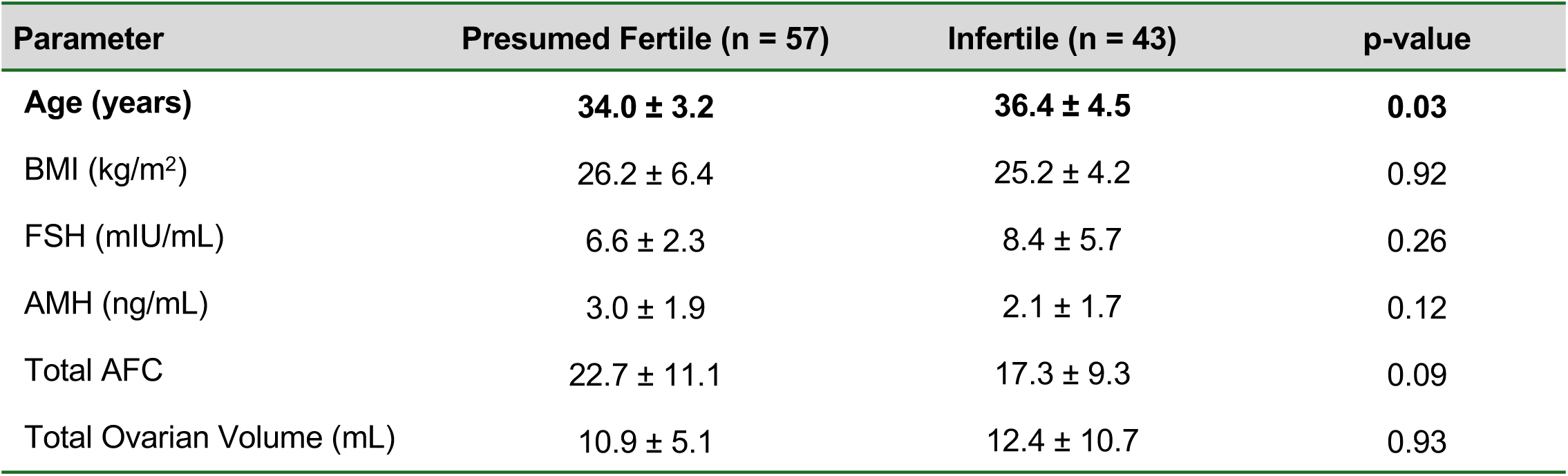
Demographic data for participants in Cohort 2 stratified by fertility status. Participants were categorized as “presumed fertile” or “infertile” based on their clinical history and reasons for seeking fertility counseling (Methods). Fertility status analysis by unpaired t-test.

## REFERENCES

1 Duncan, F. E., Confino, R. & Pavone, M. E. in Conn’s Handbook of Models for Human Aging 109–130 (Elsevier, 2018).

2 Hughes, L., Duncan, F. E. & Babayev, E. Ovarian aging: a missing diagnosis in reproductive medicine. Nature Medicine, 1–3 (2025).

3 Mishra, G. D. et al. Optimising health after early menopause. Lancet 403, 958–968 (2024). 10.1016/s0140-6736(23)02800-3

4 Davis, S. R., Pinkerton, J., Santoro, N. & Simoncini, T. Menopause—Biology, consequences, supportive care, and therapeutic options. Cell 186, 4038–4058 (2023). 10.1016/j.cell.2023.08.016

5 Jones, A. R. et al. Bone health in women with premature ovarian insufficiency/early menopause: a 23-year longitudinal analysis. Human Reproduction 39, 1013–1022 (2024). 10.1093/humrep/deae037

6 Shieh, A. et al. Associations of Age at Menopause With Postmenopausal Bone Mineral Density and Fracture Risk in Women. J Clin Endocrinol Metab 107, e561–e569 (2022). 10.1210/clinem/dgab690

7 Anagnostis, P. et al. Association between age at menopause and fracture risk: a systematic review and meta-analysis. Endocrine 63, 213–224 (2019). 10.1007/s12020-018-1746-6

8 Honigberg, M. C. et al. Association of Premature Natural and Surgical Menopause With Incident Cardiovascular Disease. JAMA 322, 2411–2421 (2019). 10.1001/jama.2019.19191

9 Muka, T. et al. Association of Age at Onset of Menopause and Time Since Onset of Menopause With Cardiovascular Outcomes, Intermediate Vascular Traits, and All-Cause Mortality: A Systematic Review and Meta-analysis. JAMA Cardiology 1, 767–776 (2016). 10.1001/jamacardio.2016.2415

10 Hao, W. et al. Age at menopause and all-cause and cause-specific dementia: a prospective analysis of the UK Biobank cohort. Hum Reprod 38, 1746–1754 (2023). 10.1093/humrep/dead130

11 Xing, Z. & Kirby, R. S. Age at natural or surgical menopause, all-cause mortality, and lifespan among postmenopausal women in the United States. Menopause 31, 176–185 (2024). 10.1097/gme.0000000000002314

12 Wu, L. et al. Macrophage iron dyshomeostasis promotes aging-related renal fibrosis. Aging Cell 23, e14275 (2024). 10.1111/acel.14275

13 Tong, C., Xue, Y., Wang, W. & Chen, X. Advanced liver fibrosis, but not MASLD, is associated with accelerated biological aging: a population-based study. BMC Public Health 24, 3293 (2024). 10.1186/s12889-024-20808-y

14 Raslan, A. A. et al. Lung injury-induced activated endothelial cell states persist in aging-associated progressive fibrosis. Nat Commun 15, 5449 (2024). 10.1038/s41467-024-49545-x

15 Schelbert, E. B. et al. Myocardial Fibrosis Quantified by Extracellular Volume Is Associated With Subsequent Hospitalization for Heart Failure, Death, or Both Across the Spectrum of Ejection Fraction and Heart Failure Stage. J Am Heart Assoc 4 (2015). 10.1161/jaha.115.002613

16 Martin, P., Pardo-Pastor, C., Jenkins, R. G. & Rosenblatt, J. Imperfect wound healing sets the stage for chronic diseases. Science 386, eadp2974 (2024). 10.1126/science.adp2974

17 Foley, K. G., Pritchard, M. T. & Duncan, F. E. Macrophage-derived multinucleated giant cells: hallmarks of the aging ovary. Reproduction 161, V5–V9 (2021).

18 Amargant, F. et al. Ovarian stiffness increases with age in the mammalian ovary and depends on collagen and hyaluronan matrices. Aging Cell 19, e13259 (2020). 10.1111/acel.13259

19 Dipali, S. S. et al. Proteomic quantification of native and ECM-enriched mouse ovaries reveals an age-dependent fibro-inflammatory signature. Aging (Albany NY) 15, 10821–10855 (2023). 10.18632/aging.205190

20 Briley, S. M. et al. Reproductive age-associated fibrosis in the stroma of the mammalian ovary. Reproduction 152, 245–260 (2016). 10.1530/REP-16-0129

21 Tang, M., Zhao, M. & Shi, Y. New insight into the role of macrophages in ovarian function and ovarian aging. Front Endocrinol (Lausanne) 14, 1282658 (2023). 10.3389/fendo.2023.1282658

22 Mara, J. N. et al. Ovulation and ovarian wound healing are impaired with advanced reproductive age. Aging (Albany NY) 12, 9686–9713 (2020). 10.18632/aging.103237

23 Landry, D. A., Vaishnav, H. T. & Vanderhyden, B. C. The significance of ovarian fibrosis. Oncotarget 11, 4366–4370 (2020). 10.18632/oncotarget.27822

24 Umehara, T., Richards, J. S. & Shimada, M. The stromal fibrosis in aging ovary. Aging (Albany NY) 10, 9–10 (2018). 10.18632/aging.101370

25 McCloskey, C. W. et al. Metformin Abrogates Age-Associated Ovarian Fibrosis. Clin Cancer Res 26, 632–642 (2020). 10.1158/1078-0432.CCR-19-0603

26 Landry, D. A. et al. Metformin prevents age-associated ovarian fibrosis by modulating the immune landscape in female mice. Science Advances 8, eabq1475 (2022). doi:10.1126/sciadv.abq1475

27 Duncan, F. E. & Babayev, E. The ovarian stroma as a therapeutic target. Science 391, 552–553 (2026). doi:10.1126/science.aee7270

28 Watson, M. A. et al. Senescence-Linked Fibrosis in the Aging Human Ovary Revealed by p16-Based Histological Profiling and Spatial Transcriptomics. bioRxiv, 2025.2012.2003.692228 (2025). 10.64898/2025.12.03.692228

29 Pietroforte, S., Plough, M. & Amargant, F. Age-associated increased stiffness of the ovarian microenvironment impairs follicle development and oocyte quality and rapidly alters follicle gene expression. bioRxiv, 2024.2006. 2009.598134 (2024).

30 Amargant, F., Magalhaes, C., Pritchard, M. T. & Duncan, F. E. Systemic low-dose anti-fibrotic treatment attenuates ovarian aging in the mouse. GeroScience, 1–21 (2024).

31 Winstanley, Y. E. et al. Emerging therapeutic strategies to mitigate female and male reproductive aging. Nature Aging 4, 1682–1696 (2024).

32 Umehara, T. et al. Female reproductive life span is extended by targeted removal of fibrotic collagen from the mouse ovary. Science Advances 8, eabn4564 (2022).

33 Zhou, F., Shi, L.-B., Zhang, S.-Y. & Ji, Y.-Y. Ovarian Fibrosis: A Phenomenon of Concern. Chinese Medical Journal 130, 365–371 (2017). doi:10.4103/0366-6999.198931

34 West, E. R., Xu, M., Woodruff, T. K. & Shea, L. D. Physical properties of alginate hydrogels and their effects on in vitro follicle development. Biomaterials 28, 4439–4448 (2007).

35 Ouni, E. et al. Spatiotemporal changes in mechanical matrisome components of the human ovary from prepuberty to menopause. Hum Reprod 35, 1391–1410 (2020). 10.1093/humrep/deaa100

36 Engel, B. et al. Quantification of extracellular matrix remodeling for the non-invasive identification of graft fibrosis after liver transplantation. Scientific Reports 13, 6103 (2023). 10.1038/s41598-023-33100-7

37 Rasmussen, D. G. K. et al. Collagen turnover profiles in chronic kidney disease. Scientific Reports 9, 16062 (2019). 10.1038/s41598-019-51905-3

38 Huang, Y. et al. Shear wave elastography of breast lesions: quantitative analysis of elastic heterogeneity improves diagnostic performance. Ultrasound in Medicine & Biology 45, 1909–1917 (2019).

39 Garra, B. S. Elastography: history, principles, and technique comparison. Abdom Imaging 40, 680–697 (2015). 10.1007/s00261-014-0305-8

40 Berg, W. A. et al. Quantitative Maximum Shear-Wave Stiffness of Breast Masses as a Predictor of Histopathologic Severity. American Journal of Roentgenology 205, 448–455 (2015). 10.2214/ajr.14.13448

41 Zaniker, E. J. et al. Shear wave elastography to assess stiffness of the human ovary and other reproductive tissues across the reproductive lifespan in health and disease. Biology of reproduction 110, 1100–1114 (2024).

42 Sigrist, R. M. S., Liau, J., Kaffas, A. E., Chammas, M. C. & Willmann, J. K. Ultrasound Elastography: Review of Techniques and Clinical Applications. Theranostics 7, 1303–1329 (2017). 10.7150/thno.18650

43 Cedars, M. I. Evaluation of female fertility—AMH and ovarian reserve testing. The Journal of Clinical Endocrinology & Metabolism 107, 1510–1519 (2022).

44 Bressler, L. H. & Steiner, A. Anti-Müllerian hormone as a predictor of reproductive potential. Current Opinion in Endocrinology, Diabetes and Obesity 25, 385–390 (2018).

45 Di Clemente, N., Racine, C., Pierre, A. & Taieb, J. Anti-Müllerian hormone in female reproduction. Endocrine reviews 42, 753–782 (2021).

46 Fong, S. L. et al. Anti-Müllerian hormone: a marker for oocyte quantity, oocyte quality and embryo quality? Reproductive biomedicine online 16, 664–670 (2008).

47 Pietroforte, S. & Amargant, F. Increased stiffness mimicking ovarian aging induces a fibroinflammatory response in follicles and impairs oocyte quality. Reproduction 171 (2026). 10.1093/reprod/xaaf026

48 Brito, I. R. et al. Alginate hydrogel matrix stiffness influences the in vitro development of caprine preantral follicles. Mol Reprod Dev 81, 636–645 (2014). 10.1002/mrd.22330

49 Wu, M. et al. Modulating IL-11-dependent matrix stiffness to delay ovarian aging. Nature Aging 6, 1395–1416 (2026). 10.1038/s43587-026-01159-2

50 Genro, V. K., Matte, U., De Conto, E., Cunha-Filho, J. S. & Fanchin, R. Frequent polymorphisms of FSH receptor do not influence antral follicle responsiveness to follicle-stimulating hormone administration as assessed by the Follicular Output RaTe (FORT). Journal of assisted reproduction and genetics 29, 657–663 (2012).

51 Grynberg, M. & Labrosse, J. Understanding follicular output rate (FORT) and its implications for POSEIDON criteria. Frontiers in endocrinology 10, 246 (2019).

52 Alviggi, C. et al. Understanding ovarian hypo-response to exogenous gonadotropin in ovarian stimulation and its new proposed marker—the follicle-to-oocyte (FOI) index. Frontiers in endocrinology 9, 589 (2018).

53 Conforti, A. et al. Management of women with an unexpected low ovarian response to gonadotropin. Frontiers in endocrinology 10, 387 (2019).

54 Hansen, N. U. B. et al. Type VIII collagen is elevated in diseases associated with angiogenesis and vascular remodeling. Clinical biochemistry 49, 903–908 (2016).

55 Dipali, S. S. et al. Proteomic quantification of native and ECM-enriched mouse ovaries reveals an age-dependent fibro-inflammatory signature. Aging 15, 10821–10855 (2023).

56 Ouni, E. et al. A blueprint of the topology and mechanics of the human ovary for next-generation bioengineering and diagnosis. Nat Commun 12, 5603 (2021). 10.1038/s41467-021-25934-4

57 Choi, Y. et al. A single-cell gene expression atlas of human follicular aspirates: Identification of leukocyte subpopulations and their paracrine factors. Faseb j 37, e22843 (2023). 10.1096/fj.202201746RR

58 Roos, K. et al. Single-cell RNA-seq analysis and cell-cluster deconvolution of the human preovulatory follicular fluid cells provide insights into the pathophysiology of ovarian hyporesponse. Frontiers in Endocrinology Volume 13 - 2022 (2022). 10.3389/fendo.2022.945347

59 Jones, A. S. K. et al. Cellular atlas of the human ovary using morphologically guided spatial transcriptomics and single-cell sequencing. Science Advances 10, eadm7506 (2024). doi:10.1126/sciadv.adm7506

60 Wu, H. et al. Single-Cell Sequencing Reveals an Intrinsic Heterogeneity of the Preovulatory Follicular Microenvironment. Biomolecules 12 (2022). 10.3390/biom12020231

61 Dipali, S. S. et al. Self-Organizing Ovarian Somatic Organoids Preserve Cellular Heterogeneity and Reveal Cellular Contributions to Ovarian Aging. Aging Cell 25, e70333 (2026). 10.1111/acel.70333

62 Fan, X. et al. Single-cell reconstruction of follicular remodeling in the human adult ovary. Nat Commun 10, 3164 (2019). 10.1038/s41467-019-11036-9

63 Meng, X. M., Nikolic-Paterson, D. J. & Lan, H. Y. TGF-β: the master regulator of fibrosis. Nat Rev Nephrol 12, 325–338 (2016). 10.1038/nrneph.2016.48

64 Isola, J. V. V. et al. A single-cell atlas of the aging mouse ovary. Nature Aging 4, 145–162 (2024). 10.1038/s43587-023-00552-5

65 Jiang, C. et al. Serpine 1 induces alveolar type II cell senescence through activating p53-p21-Rb pathway in fibrotic lung disease. Aging Cell 16, 1114–1124 (2017). 10.1111/acel.12643

66 Naba, A. et al. The matrisome: in silico definition and in vivo characterization by proteomics of normal and tumor extracellular matrices. Mol Cell Proteomics 11, M111.014647 (2012). 10.1074/mcp.M111.014647

67 Schilling, B. et al. Senescence-Linked Fibrosis in the Aging Human Ovary Revealed by p16-Based Histological Profiling and Spatial Transcriptomics. Res Sq (2026). 10.21203/rs.3.rs-8290960/v1

68 Edepli, B. G. & Yaba, A. Molecular mechanisms of ovarian fibrosis. Mol Hum Reprod 32 (2026). 10.1093/molehr/gaaf058

69 Pedrycz-Wieczorska, A. et al. CD44 as a Central Integrator of Inflammation and Fibrosis: From Molecular Signaling to Environmental Modulation. International Journal of Molecular Sciences 26, 8870 (2025).

70 Ying, H. Z. et al. PDGF signaling pathway in hepatic fibrosis pathogenesis and therapeutics (Review). Mol Med Rep 16, 7879–7889 (2017). 10.3892/mmr.2017.7641

71 Zhu, Y. et al. Ovarian remodeling and aging-related chronic inflammation and fibrosis in the mammalian ovary. J Ovarian Res 18, 133 (2025). 10.1186/s13048-025-01715-1

72 Zaniker, E. J. et al. Shear wave elastography to assess stiffness of the human ovary and other reproductive tissues across the reproductive lifespan in health and disease†. Biol Reprod 110, 1100–1114 (2024). 10.1093/biolre/ioae050

73 Sibinga, N. E. et al. Collagen VIII is expressed by vascular smooth muscle cells in response to vascular injury. Circ Res 80, 532–541 (1997). 10.1161/01.res.80.4.532

74 Li, Q. et al. COL8A1 regulates endothelial phenotype in inflammatory endothelial-to-mesenchymal transition. Am J Physiol Heart Circ Physiol 329, H1331–h1346 (2025). 10.1152/ajpheart.00339.2025

75 Shi, H., Yu, Y., Guo, K. & He, R. Type VIII collagen: advances in matrix biology and translational promise. Front Bioeng Biotechnol 13, 1732988 (2025). 10.3389/fbioe.2025.1732988

76 Lopes, J. et al. Type VIII collagen mediates vessel wall remodeling after arterial injury and fibrous cap formation in atherosclerosis. Am J Pathol 182, 2241–2253 (2013). 10.1016/j.ajpath.2013.02.011

77 Sutmuller, M., Bruijn, J. A. & de Heer, E. Collagen types VIII and X, two non-fibrillar, short-chain collagens. Structure homologies, functions and involvement in pathology. Histol Histopathol 12, 557–566 (1997).

78 Vo, N. D. N., Gaßler, N., Wolf, G. & Loeffler, I. The Role of Collagen VIII in the Aging Mouse Kidney. Int J Mol Sci 25 (2024). 10.3390/ijms25094805

79 Willumsen, N., Jorgensen, L. N. & Karsdal, M. A. Vastatin (the NC1 domain of human type VIII collagen a1 chain) is linked to stromal reactivity and elevated in serum from patients with colorectal cancer. Cancer Biol Ther 20, 692–699 (2019). 10.1080/15384047.2018.1550571

80 Williams, L., Layton, T., Yang, N., Feldmann, M. & Nanchahal, J. Collagen VI as a driver and disease biomarker in human fibrosis. Febs j 289, 3603–3629 (2022). 10.1111/febs.16039

81 Holm Nielsen, S., et al. The novel collagen matrikine, endotrophin, is associated with mortality and cardiovascular events in patients with atherosclerosis. J Intern Med 290, 179–189 (2021). 10.1111/joim.13253

82 Karsdal, M. A. et al. Assessment of liver fibrosis progression and regression by a serological collagen turnover profile. Am J Physiol Gastrointest Liver Physiol 316, G25–g31 (2019). 10.1152/ajpgi.00158.2018

83 Pilemann-Lyberg, S. et al. Markers of Collagen Formation and Degradation Reflect Renal Function and Predict Adverse Outcomes in Patients With Type 1 Diabetes. Diabetes Care 42, 1760–1768 (2019). 10.2337/dc18-2599

84 Sparding, N. et al. Endotrophin, a collagen type VI-derived matrikine, reflects the degree of renal fibrosis in patients with IgA nephropathy and in patients with ANCA-associated vasculitis. Nephrol Dial Transplant 37, 1099–1108 (2022). 10.1093/ndt/gfab163

85 Klingberg, F. et al. The fibronectin ED-A domain enhances recruitment of latent TGF-β-binding protein-1 to the fibroblast matrix. J Cell Sci 131 (2018). 10.1242/jcs.201293

86 Ma, W. et al. Direct Extracellular Matrix Modulation Attenuates Intestinal Fibrosis via a Fibronectin-Targeted Approach. Adv Sci (Weinh) 13, e19433 (2026). 10.1002/advs.202519433

87 Beyatlı, D., Brown, M., Romanescu, G. R. & Ahn, S. Aligned Fibronectin Microenvironment Temporally Facilitates Profibrotic Fibroblast Activation via Integrin α5β1. Adv Sci (Weinh), e00047 (2026). 10.1002/advs.202600047

88 Rana, T. et al. PAI-1 Regulation of TGF-β1-induced Alveolar Type II Cell Senescence, SASP Secretion, and SASP-mediated Activation of Alveolar Macrophages. Am J Respir Cell Mol Biol 62, 319–330 (2020). 10.1165/rcmb.2019-0071OC

89 Alessio, N. et al. IGFBP5 is released by senescent cells and is internalized by healthy cells, promoting their senescence through interaction with retinoic receptors. Cell Commun Signal 22, 122 (2024). 10.1186/s12964-024-01469-1

90 Zhao, T. et al. Platelet-derived growth factor-D promotes fibrogenesis of cardiac fibroblasts. Am J Physiol Heart Circ Physiol 304, H1719–1726 (2013). 10.1152/ajpheart.00130.2013

91 Klinkhammer, B. M., Floege, J. & Boor, P. PDGF in organ fibrosis. Mol Aspects Med 62, 44–62 (2018). 10.1016/j.mam.2017.11.008

92 Buhl, E. M. et al. The role of PDGF-D in healthy and fibrotic kidneys. Kidney Int 89, 848–861 (2016). 10.1016/j.kint.2015.12.037

93 Acharya, P. S. et al. Fibroblast migration is mediated by CD44-dependent TGF beta activation. J Cell Sci 121, 1393–1402 (2008). 10.1242/jcs.021683

94 Govindaraju, P., Todd, L., Shetye, S., Monslow, J. & Puré, E. CD44-dependent inflammation, fibrogenesis, and collagenolysis regulates extracellular matrix remodeling and tensile strength during cutaneous wound healing. Matrix Biol 75-76, 314-330 (2019). 10.1016/j.matbio.2018.06.004

95 Isola, J. V. V. et al. Canagliflozin treatment prevents follicular exhaustion and attenuates hallmarks of ovarian aging in genetically heterogenous mice. Geroscience 47, 3061–3076 (2025). 10.1007/s11357-024-01465-w

96 Lin, Z. et al. Antifibrotic drug finerenone restores fertility in premature ovarian insufficiency. Science 391, eadz4075 (2026). doi:10.1126/science.adz4075

97 Li, P. & Chen, Z. Association of follicle-to-oocyte index and clinical pregnancy in IVF treatment: A retrospective study of 4,323 fresh embryo transfer cycles. Front Endocrinol (Lausanne) 13, 973544 (2022). 10.3389/fendo.2022.973544

98 Chen, L. et al. Follicular Output Rate and Follicle-to-Oocyte Index of Low Prognosis Patients According to POSEIDON Criteria: A Retrospective Cohort Study of 32,128 Treatment Cycles. Front Endocrinol (Lausanne) 11, 181 (2020). 10.3389/fendo.2020.00181

99 Curchoe, C. L., Tarafdar, O., Aquilina, M. C. & Seifer, D. B. SART CORS IVF registry: looking to the past to shape future perspectives. Journal of assisted reproduction and genetics 39, 2607–2616 (2022).

100 Gokyer, D. et al. Collection of Human Follicular Fluid, Follicle Somatic Cells, and Immature Oocytes from Individuals Undergoing In Vitro Fertilization. J Vis Exp (2025). 10.3791/69122

101 Petrov, P. B., Considine, J. M., Izzi, V. & Naba, A. Matrisome AnalyzeR – a suite of tools to annotate and quantify ECM molecules in big datasets across organisms. Journal of Cell Science 136 (2023). 10.1242/jcs.261255

102 Coppé, J. P., Desprez, P. Y., Krtolica, A. & Campisi, J. The senescence-associated secretory phenotype: the dark side of tumor suppression. Annu Rev Pathol 5, 99–118 (2010). 10.1146/annurev-pathol-121808-102144

103 Zhou, Y. et al. Metascape provides a biologist-oriented resource for the analysis of systems-level datasets. Nat Commun 10, 1523 (2019). 10.1038/s41467-019-09234-6

104 Xie, Z. et al. Gene Set Knowledge Discovery with Enrichr. Curr Protoc 1, e90 (2021). 10.1002/cpz1.90

105 Kuleshov, M. V. et al. Enrichr: a comprehensive gene set enrichment analysis web server 2016 update. Nucleic Acids Res 44, W90–97 (2016). 10.1093/nar/gkw377

106 Chen, E. Y. et al. Enrichr: interactive and collaborative HTML5 gene list enrichment analysis tool. BMC Bioinformatics 14, 128 (2013). 10.1186/1471-2105-14-128

107 Jin, S. et al. Inference and analysis of cell-cell communication using CellChat. Nature Communications 12, 1088 (2021). 10.1038/s41467-021-21246-9

